# Template switching by a group II intron reverse transcriptase: biochemical analysis and implications for RNA-seq

**DOI:** 10.1101/792986

**Authors:** Alfred M. Lentzsch, Jun Yao, Rick Russell, Alan M. Lambowitz

## Abstract

The reverse transcriptases (RTs) encoded by mobile group II intron and other non-LTR-retro-elements differ from retroviral RTs in being able to template switch from the 5’ end of one template to the 3’ end of another without pre-existing complementarity between the donor and acceptor nucleic acids. Here, we used the ability of a thermostable group II intron RT (TGIRT; GsI-IIC RT) to template switch directly from synthetic RNA template/DNA primer duplexes having either a blunt end or a 3’-DNA overhang end to establish a complete kinetic framework for the reaction and identify conditions that more efficiently capture acceptor RNAs or DNAs. The rate and amplitude of template switching are optimal from starter duplexes with a single nucleotide 3’-DNA overhang complementary to the 3’ nucleotide of the acceptor RNA, suggesting a role for non-templated nucleotide addition of a complementary nucleotide to the 3’ end of cDNAs synthesized from natural templates. Longer 3’-DNA overhangs progressively decrease the rate of template switching, even when complementary to the 3’ end of the acceptor template. The reliance on a single base pair with the 3’ nucleotide of the acceptor together with discrimination against mismatches and the high processivity of the enzyme enable synthesis of full-length DNA copies of nucleic acids beginning directly at their 3’ end. We discuss possible biological functions of the template-switching activity of group II intron and other non-LTR-retroelements RTs, as well as the optimization of this activity for adapter addition in RNA-and DNA-seq.

## Introduction

Reverse transcriptases (RTs) typically have the ability to template switch during cDNA synthesis, thereby joining discontinuous nucleic acid sequences. Such template switching plays a critical role in the life cycle of retroviruses and other retroelements and also has biotechnological applications, particularly in RNA-seq, where it can serve as an alternative to RNA tailing or ligation for adding RNA-seq adapters to target RNAs (1–9). Although studied in greatest detail for retroviral RTs, the mechanism of template switching differs for group II intron and other non-long terminal repeat (non-LTR) retroelement RTs in ways that are only just beginning to be fully appreciated and exploited for biotechnological applications.

Template switching was discovered in early studies of retroviral replication, which revealed an unexpected but crucial role for two template switches, also referred to as strand transfers, for the synthesis of full-length viral genomes (1–3). Both template switches occur when the RT reaches the end of one of the LTRs of the viral RNA and involve base pairing of a singlestranded cDNA, exposed by RNase H digestion of the RNA template, to a complementary sequence in the other LTR. Template switches between internal regions of the viral RNA are also common (10). Because retroviruses package two, often nonidentical, genomic RNAs and their RTs dissociate readily from RNA templates, template switching to complementary regions of another genomic RNA occur frequently (3–12 times per replication cycle), leading to high rates of recombination between viral genomes (11, 12). Such recombination of genomic RNA sequences provides a means of rapidly propagating beneficial mutations that help retroviruses evade host defenses (13–15). Template switching by a retroviral RT is also employed for RNA-seq adapter addition in a method called SMART-Seq (Switching Mechanism At the 5’ end of the RNA Transcript sequencing), which utilizes the ability of Moloney murine leukemia virus RT to add non-templated C residues to the 3’ end of a completed cDNA, which can then base pair and enable template switching to the 3’ end of an adapter DNA oligonucleotide ending with G residues (4, 7, 16). Biochemical studies showed that efficient template switching by HIV-1 RT requires at least two base pairs between the 3’ end of the cDNA and the acceptor template, one of which must be a GC or CG base pair (17, 18).

Unlike retroviral RTs, which rely on relatively stable base-pairing interactions for template switching, the RTs encoded by non-LTR-retroelement RTs have the ability to template switch from the 5’ end of a donor template to the 3’ end of an acceptor template with little or no pre-existing complementarity between the cDNA and the acceptor (5). Non-LTR-retroelements are a broad class that includes non-LTR-retrotransposons, such as human long interspersed nuclear element 1 (LINE-1) and insect R2 retroelements, as well as mitochondrial retroplasmids (MRPs), and bacterial and organellar mobile group II introns, which are evolutionary ancestors of non-LTR-retrotransposons and retroviruses in eukaryotes (19, 20). The RTs of non-LTR-retroelements differ structurally from retroviral RTs, with distinctive features not found in retroviral RTs including an N-terminal extension (NTE) containing a conserved sequence block (RT0) and two structurally conserved insertions (RT2a and 3a) between the universally conserved RT sequence blocks (RT1-7) (21).

Biochemical studies of template switching by three non-LTR-retroelement RTs have revealed common features as well as differences that may reflect adaptations of this activity for the life cycle of different retroelements. The Mauriceville and Varkud MRPs found in some *Neurospora* spp. strains are closely related small closed-circular DNAs, which encode an RT that functions in plasmid replication (22, 23). The MRPs are transcribed to yield full-length transcripts with a 3’-tRNA-like structure and 3’ CCA, which are recognized by the RT for the initiation of cDNA synthesis at the penultimate C of the 3’ CCA either *de novo* (*i.e.*, without a primer) or by using the 3’ end of either a DNA or RNA primer with little or no complementarity to the RNA template (24, 25). Upon reaching the 5’ end of the plasmid transcript, template switching to the 3’ end of the same or another plasmid transcript yields multimeric cDNAs, which can undergo intramolecular recombination to regenerate closed-circular plasmid DNA (25–29). End-to-end template switching also occurs between the MRP transcript and heterologous mtDNA transcripts, leading to recombinant plasmids containing an integrated mtDNA sequence, frequently a mt tRNA (27, 30, 31). Such recombinant plasmids have a replicative advantage over the wild-type plasmid, as well as the ability to stably integrate into mtDNA by homologous recombination at the corresponding mtDNA locus without relying on a specialized DNA integrase (27, 30, 31).

The insect R2 element is a non-LTR-retrotransposon that inserts site-specifically into the large subunit rRNA gene by a target-primed reverse transcription (TPRT) mechanism in which the RT nicks one strand of the rDNA and then uses the 3’ end of the cleaved DNA strand as a primer to synthesize a full-length cDNA beginning at the 3’ end of the R2 RNA (32). Biochemical studies showed that upon reaching the 5’ end of an RNA template, the RT can template switch to the 3’ end of another RNA or DNA template and that this template switching is facilitated by non-templated nucleotide addition (NTA) of short stretches of complementary nucleotides (“microhomologies”) to the 3’ end of the completed cDNA (33, 34). Such template switching, utilizing microhomologies generated by NTA, has been proposed to play a role in linking the 5’ end of the R2 element to host genomic sequences upstream of the cleavage site and in the formation of chimeric integration products, containing additional heterologous sequences (31–33, 35–37).

Finally, mobile group II introns, the type of non-LTR-retroelement whose RT is studied here, are bacterial and organellar retrotransposons that insert site-specifically into DNA target sites by a process called retrohoming. In this process, the excised intron lariat RNA integrates directly into a DNA strand by reverse splicing and is reverse transcribed by the RT using either an opposite strand nick or a nascent DNA strand as a primer for reverse transcription (20, 38). Group II intron RTs have an end-to-end template-switching activity similar to that of MRP and R2 element RTs (6, 39), and this activity has been utilized for adapter addition in RNA-seq using a thermostable group II intron RT (TGIRT-seq) (6, 8, 9). Unlike the R2 element RT (33, 34), group II intron RTs do not need to be actively elongating to template switch and can do so directly from synthetic RNA template/DNA primer substrates with a single-nucleotide (1-nt) 3’-DNA overhang that base pairs with the 3’ nucleotide of an RNA acceptor (6). This reaction enables the precise attachment of 3’ RNA-seq adapters to target RNAs or DNAs in a single step without tailing or ligation (8, 9, 40).

Recently, we determined a crystal structure of a full-length thermostable group II intron RT (GsI-IIC RT) in complex with template-primer substrate and incoming dNTP that provided insight into the structural basis for template switching by non-LTR-retroelement RTs (41). This crystal structure revealed that the NTE and RT0 loop present in non-LTR-retroelement RTs likely form a binding pocket for the 3’ end of an acceptor nucleic acid, a structure that does not exist in retroviral RTs. Additionally, we found that mutations affecting the RT0 loop of the GsI-IIC RT, which forms the lid of the binding pocket, strongly inhibit template switching while minimally affecting primer-extension activity. A similar finding was reported for alanine substitution in the conserved RT0 motif of the R2 RT (42), suggesting that the RT0-lidded binding pocket plays a similar role in template switching in other non-LTR-retroelement RTs. The crystal structure of GsI-IIC RT opens the possibility of detailed structure-function analysis of the template-switching activity of group II intron and homologous non-LTR-retroelement RTs, as well as its optimization for RNA-seq applications.

Here, we took advantage of the ability of GsI-IIC RT to template switch directly from synthetic RNA template/DNA primer substrates to carry out a detailed kinetic analysis of the template switching and related NTA activities of GsI-IIC RT (sold commercially as TGIRT-III). We found that the rate and amplitude of template switching are optimal from starter duplexes with a single nucleotide 3’-DNA overhang complementary to the 3’ nucleotide of the acceptor RNA. We also show that this single base pair to the 3’ nucleotide of a target RNA dictates template switching with high fidelity and yields seamless junctions, while template switching from blunt-end duplexes is inefficient and yields heterogenous junctions. Our results inform possible biological functions of template switching for non-LTR-retroelement RTs and suggest how to further optimize this activity for adaptor addition in RNA-seq.

## Results

### Overview of the template-switching reaction and determination of saturating enzyme concentrations

Fig. 1A depicts the assay that we used to analyze the template-switching activity of GsI-IIC RT, which is based on the method used for 3’-adapter addition in TGIRT-seq (6, 8, 9). In this assay, GsI-IIC RT bound to an artificial RNA template/DNA primer duplex (termed “starter duplex” or “donor”) initiates cDNA synthesis by template switching to the 3’ end of an acceptor nucleic acid. For most of our experiments, the starter duplex was the same as that used for TGIRT-seq and consists of a 34-nt RNA oligonucleotide that contains an Illumina R2 adapter sequence and has a 3’ blocking group (3SpC3; IDT) annealed to a complementary 35-nt DNA primer termed R2R (Table S1). The latter leaves a 1-nt 3’ overhang (N) that can direct template switching by base pairing to the 3’ nucleotide (N’) of the acceptor nucleic acid (6). Upon the addition of dNTPs (an equimolar mix of dATP, dCTP, dGTP, and dTTP), GsI-IIC RT initiates reverse transcription by template switching to the 3’ end of the acceptor nucleic acid and synthesizes a full-length cDNA with the R2R DNA primer attached to its 5’ end. In biochemical experiments, the DNA primer was 5’ end-labeled, allowing the products to be analyzed and quantified by denaturing PAGE (Fig. 1A, bottom left). Alternatively, in some experiments, we analyzed products by RNA-seq, which involves the ligation of a 5’-adenylated R1R adapter to the 3’ end of the cDNA followed by PCR to add capture sites and indexes for Illumina sequencing (Fig. 1A, bottom right).

**FIGURE 1.**
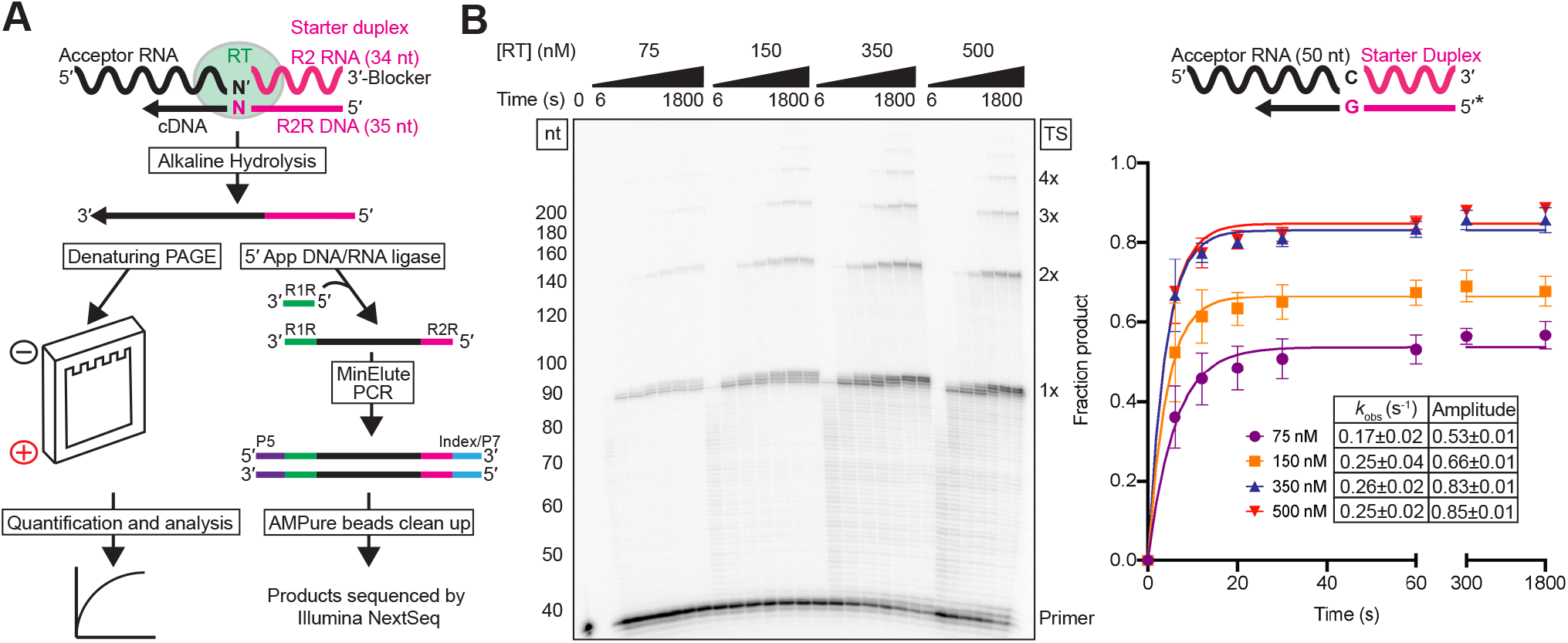
Overview of template-switching experiments and determination of saturating enzyme concentrations. *A*, Outline of the experiments. GsI-IIC RT was pre-bound to a starter duplex (magenta) consisting of a 34-nt RNA oligonucleotide containing an Illumina Read 2 (R2) sequence annealed to a complementary 35-nt DNA primer (R2R) leaving a 1-nt, 3’-DNA overhang (N) (Table S1). The 3’-overhang nucleotide (N) base pairs with the 3’ nucleotide (N’) of an acceptor RNA (black) for template switching, leading to the synthesis of a full-length cDNA copy of the acceptor RNA with the R2R adapter linked to its 5’ end. The cDNAs were incubated with NaOH to degrade RNA and neutralized with equimolar HCl prior to further analysis. For the biochemical experiments (left branch), the R2R DNA primer in the starter duplex was 5’-^32^P-labeled (*), and the cDNA products were analyzed by electrophoresis in a denaturing polyacrylamide gel, which was dried and quantified with a phosphorimager. For RNA-seq experiments (right branch), the cDNAs were cleaned up by using a MinElute column (Qiagen; not shown) prior to ligating a 5’-adenylated R1R adapter to the 3’ end of the cDNA using the thermostable 5’ app DNA/RNA ligase (New England BioLabs). After another MinElute clean-up, Illumina RNA-seq capture sites (P5 and P7) and indexes were added by PCR, and he resulting libraries were cleaned up by using AMPure XP beads (Beckman Coulter) prior to sequencing on an Illumina NextSeq 500. *B*, Determination of saturating enzyme concentrations. Template-switching reactions included various concentrations of GsI-IIC RT as indicated, 50 nM RNA template/DNA primer starter duplex (5’-^32^P-labeled on DNA primer indicated by *), 100 nM of a 50-nt RNA acceptor template, and 4 mM dNTPs (an equimolar mix of 1 mM dATP, dCTP, dGTP, and dTTP) in reaction medium containing 200 mM NaCl at 60 °C. Aliquots were quenched at times ranging from 6 to 1,800 s, and the products were analyzed by denaturing PAGE, as described in Experimental Procedures. The numbers to the left of the gel indicate size markers (a 5’ ^32^P-labeled ssDNA ladder; ss20 DNA Ladder, Simplex Sciences) run in a parallel lane, and the labels to the right of the gel indicate the products resulting from the initial template switch (1x) and subsequent end-to-end template switches from the 5’ end of one acceptor to the 3’ end of another (2x, 3x, *etc*.). The plot at the right shows time courses for the production of template-switching products (*i.e.*, products >2 nt larger than the primer), with each data set fit by a single-exponential function and the error bars indicating the standard deviations for three experiments. The inset table indicates the *k*_obs_ and amplitude parameters obtained from the fit of an exponential function to the average values from three independent determinations, along with standard errors obtained from the fit (see Experimental Procedures).

To determine an optimal enzyme concentration for the reaction, we measured the rate and amplitude of the cDNA product formation at different concentrations of GsI-IIC RT using 100 nM of a 50-nt acceptor RNA with a 3’ C residue and 50 nM of a starter duplex with a complementary 1-nt 3’ G overhang in reaction medium containing 200 mM NaCl (Fig. 1B). At all enzyme concentrations, we observed a prominent template-switching product containing the labeled DNA primer attached to a full-length cDNA of the RNA acceptor (denoted 1x), as well as a ladder of larger products resulting from sequential end-to-end template switches from the 5’ end of one acceptor RNA to the 3’ end of another acceptor RNA (denoted 2x, 3x, *etc*.; Fig. 1B). In addition to template-switching products, a fraction of the DNA primer was extended by only 1 to 3 nucleotides due to NTA to the 3’ end of the DNA primer (seen most prominently at longer time points at higher RT concentrations in Fig. 1B and analyzed further below).

Quantitation of the products resulting from template switching and extension by the RT (defined as those >2 nt larger than the primer) revealed a dominant fast phase of product formation and a minor slow phase, the latter possibly reflecting heterogeneity in the substrate and/or enzyme. For simplicity, we fit the data in this and subsequent figures using a single-exponential function to define the properties of the dominant fast phase. We found that 350-500 nM enzyme gives maximal values of the observed rate constant (~0.25 s^−1^) and reaction amplitude (~0.85) (Fig. 1B). At these enzyme concentrations, nearly all of the starter duplex (50 nM) was utilized, and excess acceptor RNA was used for multiple template switches at the longer time points. Therefore, we used 500 nM GsI-IIC RT in all subsequent reactions.

### Template switching to RNA or DNA is more efficient at lower salt concentrations

We next analyzed the effects of salt concentration on the kinetics of template switching to RNA and DNA acceptor oligonucleotides (Fig. 2). The observed rate constants for template switching to a 50-nt acceptor RNA or DNA were the same within error and relatively unaffected by NaCl concentrations between 100 and 300 mM, but decreased 2- to 3-fold at 400 mM NaCl (Fig. 2). The maximum amplitude of product formation was somewhat higher for the RNA than the DNA acceptor (0.85 and 0.77, respectively at 200 mM NaCl), but in both cases decreased by ~40% at 400 mM NaCl, as did the proportion of products resulting from multiple template switches (Fig. 2). Additional experiments showed that neither the acceptor RNA nor the enzyme became limiting at 400 mM NaCl (Fig. S1), suggesting that the high salt concentration may adversely affect the conformation of the enzyme or substrates for template switching. By contrast, in a parallel primer extension reaction in which GsI-IIC RT initiated synthesis from a DNA primer annealed to the 3’ end of a 1.1-kb RNA template, the rate of extension decreased from ~9 nt/s at 100 mM NaCl to ~4 nt/s at 400 mM NaCl but reached the same amplitude irrespective of salt concentration (Fig. S2). These findings indicate that the decreased amplitude of reverse transcription reactions that require template switching results from decreased efficiency of the initial template switch and not the subsequent reverse transcription. A reciprocal experiment using non-denaturing gel electrophoresis to examine utilization of a 5’-^32^P-labeled acceptor RNA showed that a higher proportion of the acceptor was utilized for template-switching at 200 mM than at 450 mM NaCl, a salt concentration used in TGIRT-seq to suppress NTA and multiple template switches (63% and 19% of acceptor utilized at 200 and 450 mM NaCl, respectively; Fig. S3). Based on these results, we used 200 mM NaCl in most subsequent experiments.

**FIGURE 2.**
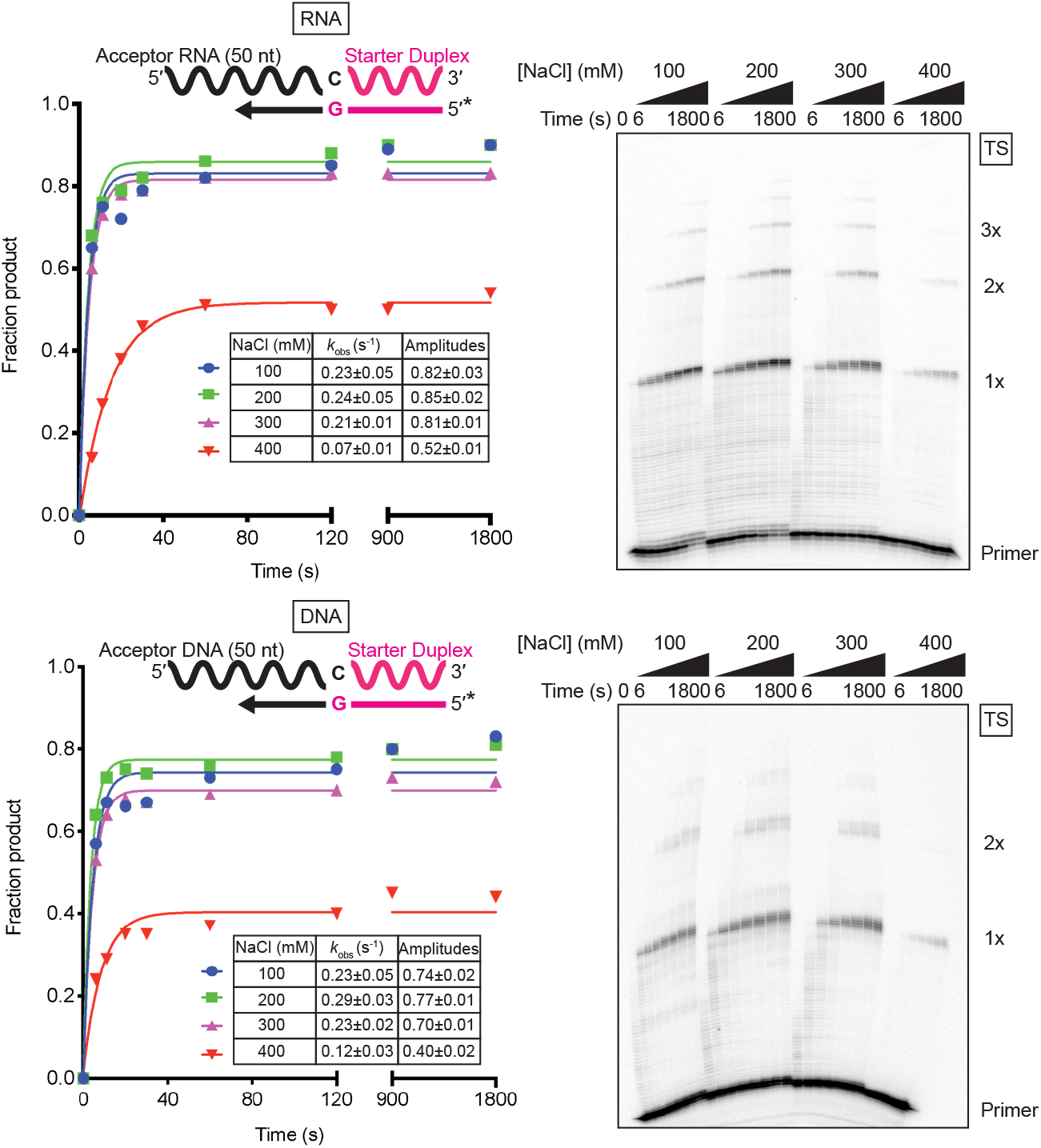
Template switching to RNA and DNA is more efficient at lower salt concentrations. Template-switching reaction time courses with 100 nM of 50-nt RNA (top) or DNA (bottom) acceptors of identical sequence (Table S1) were done with 500 nM GsI-IIC RT and 50 nM starter duplex with a 5’ ^32^P-labeled (*) DNA primer in reaction medium containing 100, 200, 300, or 400 mM NaCl. Time points were taken at intervals ranging from 6 to 1,800 s and the products were analyzed by denaturing PAGE, as described in Fig. 1B. The plots to the left of the gel show the data fit by a single-exponential function to calculate the *k*_obs_ and amplitude for each time course, and the values are summarized in the inset tables together with the standard error of the fit. The gels are labeled as in Fig. 1.

### Lower salt concentration increases template switching to RNAs ending with 3’-phosphate or 2’-O-Me groups

Previous work showed that in a reaction medium containing 450 mM NaCl, template switching of group II intron RTs is most efficient to RNAs with a 3’-OH group and strongly inhibited for RNAs with either a 3’-phosphate or 2’-*O*-Me group (6). In light of our finding that template switching is more efficient at lower salt concentrations, we assayed the template-switching activity of GsI-IIC RT to otherwise identical 21-nt RNA acceptors with these modifications in reaction media containing 200 mM NaCl and made parallel measurements at 450 mM NaCl for comparison (Fig. 3). We found that template switching to RNAs with a 3’-phosphate or a 2’-*O*-Me group was substantially more efficient at 200 mM NaCl than at 450 mM NaCl, albeit with rates and amplitudes remaining lower than those for the same RNA with a 3’-OH group (Fig. 3). The inhibitory effects of these modifications presumably reflect that they affect the binding or alignment of the 3’ end of the acceptor nucleic acid in the template-switching pocket. We also examined template switching to DNA acceptors of identical sequence without or with a dideoxy nucleotide at their 3’ end. Template switching was not impeded by the absence of a 3’-OH group, indicating that recognition of a 2’- or 3’-OH group of the acceptor is not required for template switching.

**FIGURE 3.**
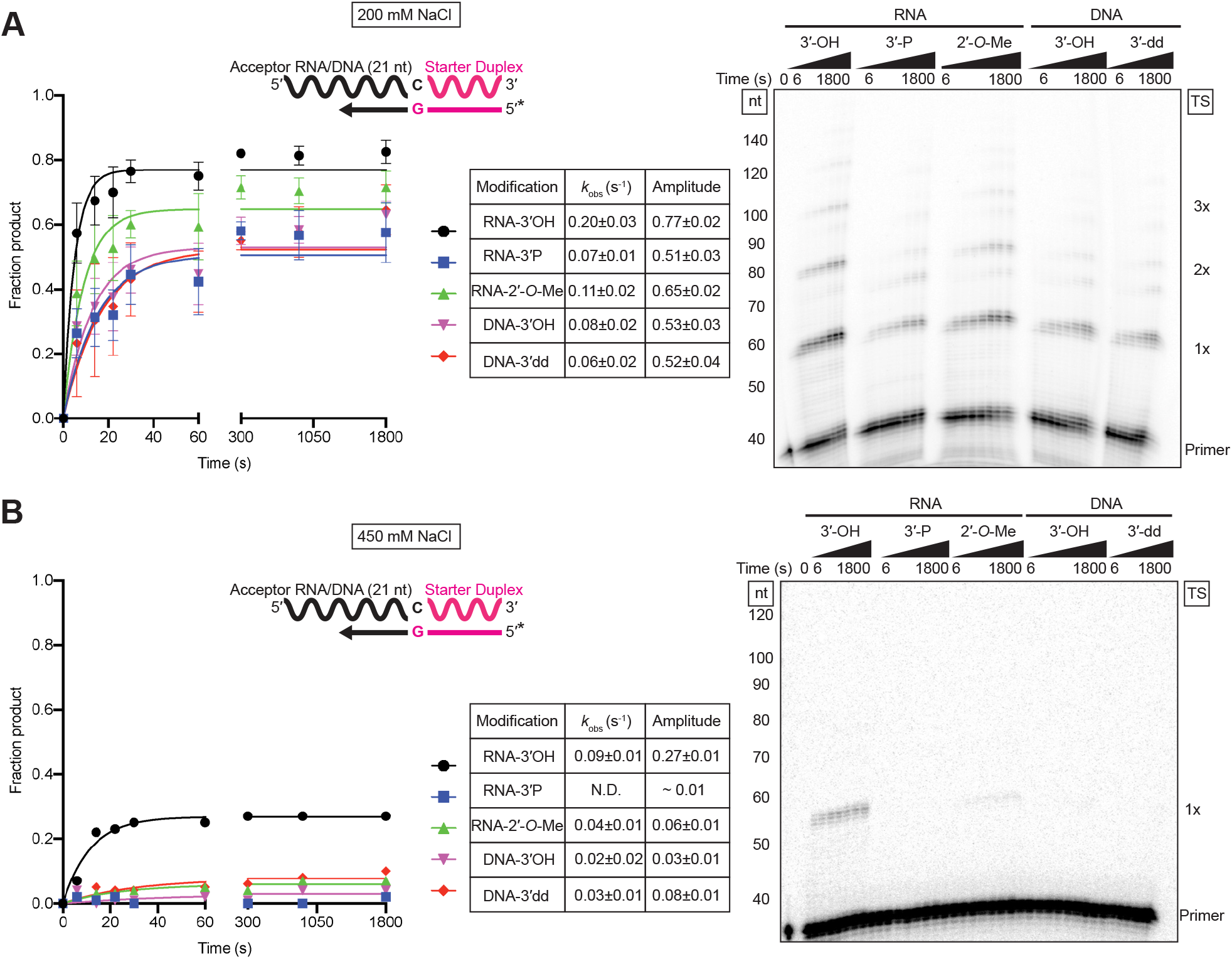
A lower salt concentration increases template switching to acceptor RNAs ending with 3’-phosphate or 2’-*O*-Me groups. *A* and *B*, Time courses of template switching to 21-nt RNA and DNA acceptors of identical sequence but different 3’-end modifications (Table S1) at 200 and 450 mM NaCl, respectively. The RNA acceptors had a 3-hydroxyl (OH), 3’-phosphate (3’ P) or 2’-*O*-methyl (2’-*O*-Me) group, and the DNA acceptors had a 3’ hydroxyl (OH) or a dideoxy (3’ dd) terminus. Reactions were done using 500 nM GsI-IIC RT, 100 nM acceptor RNA or DNA, and 50 nM starter duplex with a 5’-^32^P-lableled (*) DNA primer. Time points were taken at intervals ranging from 6 to 1,800 s, and the products were analyzed by denaturing PAGE, as described in Fig. 1. The plots to the left of the gel show the data fit by a single-exponential function to calculate the *k*_obs_ and amplitude for each reaction, and the values and standard errors of the fit are shown in the tables to the right of the plots. The gels are labeled as in Fig. 1.

### Non-templated nucleotide addition activity of GsI-IIC RT using mixed dNTPs

The ability of GsI-IIC RT to add non-templated nucleotides to the 3’ end of the DNA primer in the starter duplex could affect the efficiency of template switching positively or negatively. To probe this NTA activity quantitatively, we incubated GsI-IIC RT with a blunt-end version of the starter duplex and varying concentrations of an equimolar mixture of all four dNTPs. Gel analysis of NTA reactions showed progressive addition of up to three nucleotides (Fig. 4A). The product band reflecting addition of a single nucleotide accumulated and persisted throughout the reaction, indicating that the first nucleotide addition was faster than the second and/or that a fraction of the complex entered a paused or stopped state, a scenario discussed further below. Accordingly, the intensities of all of the product bands were summed to estimate the rate of step 1, product bands 2 and 3 were summed to estimate the rate of step 2, and product band 3 was used to estimate the rate of step 3 (Fig. 4B and Fig. S4A).

**FIGURE 4.**
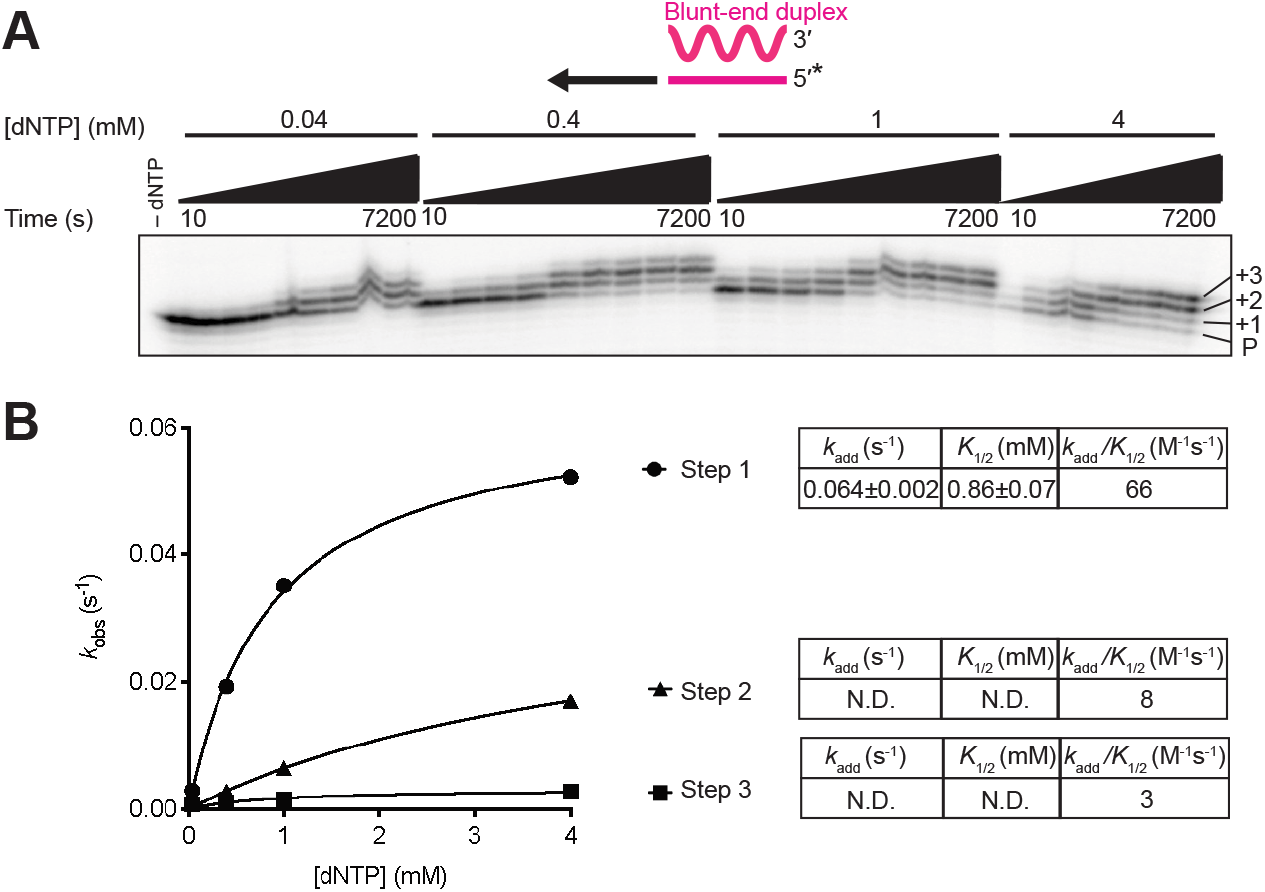
Non-templated nucleotide addition activity of GsI-IIC RT using a mixture of all four dNTPs. Reactions included 500 nM GsI-II RT and 50 nM of a blunt-end starter duplex with 5’-^32^P-labeled (*) DNA primer in reaction medium containing 200 mM NaCl and varying dNTP concentrations (0.04, 0.4, 1, and 4 mM, where 4 mM is an equimolar mix of 1 mM dATP, dCTP, dGTP, and dTTP). Aliquots were stopped after times ranging from 10 to 7,200 s, and the products were analyzed by electrophoresis in a denaturing polyacrylamide gel, which was dried and scanned with a phosphorimager. Each product band was quantified individually and summed to estimate the rate of step 1. Product bands 2 and 3 were summed to estimate the rate of step 2, and product band 3 was used to estimate the rate of step 3. The data were plotted and fit by a single-exponential function to calculate the *k*_obs_ and amplitude parameters for each reaction. *A*, Representative gel showing the labeled DNA primer (P) and bands resulting from NTA of 1, 2, and 3 nucleotides to the 3’ end of the DNA. *B*, A plot of *k*_obs_ values as a function of dNTP concentration fit by a hyperbolic function to calculate *k*_add_, the catalytic rate at saturating substrate concentration; *K*_1/2_, the substrate concentration at half maximum *k*_add_; and *k*_add_/K_1/2_ for each NTA step (values summarized in tables to the right of the plots). The individual parameter values *k*_add_ and *K*_1/2_ were not well defined for steps two and three because saturation was not reached at 4 mM dNTP, and they are therefore indicated as N.D. (not determined). Although the progress curve of the second and third NTA products (Fig. S4) would be expected to include kinetic lags in principle, the rate constants for NTA are progressively lower with repeated additions, such that the data for these additions are adequately described by simple exponential functions without lag phases. All reactions were performed at least twice, and some time points were collected three times. Data were averaged for each time point, and these averages were fit by a single-exponential function to obtain the *k*_obs_ values.

Our analysis of the rate dependence on dNTP concentration showed that the first dNTP addition was the fastest and most efficient, with a maximal rate constant (*k*_add_) of 0.064 ± 0.002 s^−1^ and a second-order rate constant (*k*_add_/*K*_1/2_) of 66 M^−1^ s^−1^ (Fig. 4B and Fig. S4A, left). The observed rate constants for the second and third NTA were lower than that for the first NTA at each dNTP concentration and did not approach a maximum at 4 mM dNTP (Fig. 4B and Fig. S4A, right), resulting in lower *k*_add_/*K*_1/2_ values of 8 M^−1^ s^−1^ and 3 M^−1^ s^−1^, respectively. Thus, each step of NTA is progressively slower, with the second step approximately 8-fold slower than the first step, and the third step even slower than the second step (Fig. 4B). We also measured NTA using a starter duplex with a 1-nt overhang (Fig. S4B). As expected, the kinetic parameters for addition of the first nucleotide from a 1-nt overhang duplex were similar to those determined for the second step starting from a blunt-end starter duplex (compare Fig. S4B to Fig. 4A and Fig. S4A). Thus, the data indicate that the first NTA occurs most efficiently, whereas the second and third additions are substantially and increasingly less efficient.

### Nucleotide preferences for non-templated nucleotide addition activity

To probe the nucleotide preferences of NTA activity, we compared NTA reactions performed with each of the four dNTPs individually. This comparison showed that the order of efficiencies of the first NTA from a blunt-end duplex was A>G>>C≈T (Fig. 5A, Fig. S5, and Table S2), indicating a preference for purines and suggesting that incorporation is largely governed by the “A-rule,” which is followed by a variety of DNA polymerases and RTs (34, 43, 44). Interestingly, for the first NTA from a starter duplex with a 3’-G overhang (the equivalent of a second NTA for a blunt-end duplex), dATP addition was even more strongly preferred (7- to 17-fold higher *k*_add_/*K*_1/2_ than for dGTP, dTTP, or dCTP; Fig. 5A).

**FIGURE 5.**
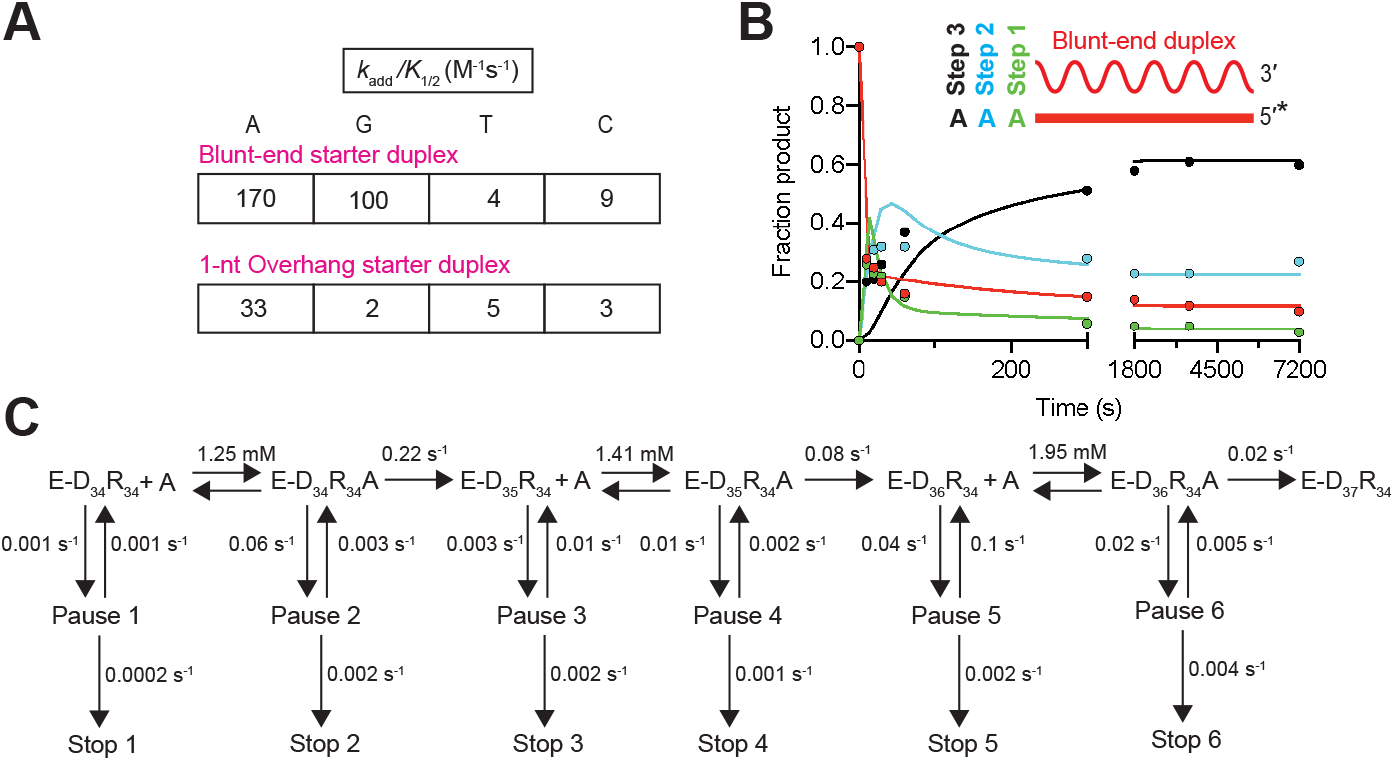
Nucleotide preferences for non-templated nucleotide addition activity. *A*, The second-order rate constants of NTA activity using individual dNTPs for the first step (Blunt-end starter duplex, top) and second step (1-nt G overhang starter duplex, bottom) are shown. The kinetic parameters for *k*_add_ and *K*_1/2_ were obtained as described in Fig. 4. Gels and plots are shown in Fig. S5, reaction conditions are given in the Fig. S5 legend, and kinetic values are summarized in Table S1. *B*, Plots showing global data fitting of consecutive dATP addition (Fig. S5) to a blunt-end starter duplex at 4 mM dATP. Each color represents a unique species in the reaction pathway: red, blunt-end starter duplex; green, product after first NTA; blue, product after second NTA; black, product after third NTA. *C*, The global model used for NTA reactions with dATP is shown with the parameters obtained from global fitting. Analogous schemes for NTA of the other dNTPs are shown in Fig. S6.

Closer analysis of the NTA progress curves revealed that even at the reaction endpoints, significant fractions of the primer do not undergo the maximum of 3-4 NTAs (*e.g.* Fig. 4A), suggesting complex kinetics with potential side reactions and/or pausing and stopping mechanisms. To explore these possibilities and to generate a model that recapitulates all features of the data, we analyzed the single dNTP reactions by global simulation and fitting (45) (Fig. 5 (B and C), Fig. S5, and Fig. S6). For each dNTP, the NTA progress curves using four concentrations of nucleotide were fit globally to reaction schemes that include up to three rapid-equilibrium nucleotide-binding steps, the second and third of which require translocation, and up to three steps of nucleotide addition (Fig. 5C and Fig. S6). Supporting and extending the conclusions from the mixed dNTP reactions (Fig. 4), for each dNTP the first addition is the most efficient and the second and third additions are progressively less efficient. Interestingly, the best-fit models also include both paused and stopped states, as suggested by the multi-phase kinetics and lack of complete decay of intermediates (Fig. 5B and Fig. S6). Paused states have been observed for translesion polymerases, with pausing occurring when the 3’ end of the primer is in a pretranslocation state occupying the nucleotide binding pocket (46, 47). The stopped state may reflect a further inhibited enzyme state that occurs during NTA or dissociation of the enzyme.

We also explored the starter duplex requirements for NTA (Fig. S7). NTA product formation was slightly higher for a blunt-end RNA/DNA starter duplex than for a blunt-end DNA/DNA starter duplex of the same nucleotide sequence. However, in contrast to terminal transferase activity (48), NTA by GsI-IIC RT did not occur efficiently to a single-stranded DNA primer under our experimental conditions (Fig. S7).

Overall, our findings show that purines are preferred for the first NTA and dATP is strongly preferred for the second NTA step. The findings that the first nucleotide addition from a blunt-end duplex is faster and more efficient than subsequent additions and that dATP is preferred for both the first and second NTAs may be relevant to the physiological functions of NTA and template switching by group II intron RTs (see Discussion).

### Template switching is favored by a single nucleotide 3’overhang

The difference in efficiency between the first and subsequent NTAs by GsI-IIC RT prompted us to investigate template-switching activity from different length 3’-DNA overhangs. To this end, we compared the template-switching activity from a blunt-end starter duplex to that from starter duplexes with 1-, 2-, or 3-nt 3’-DNA overhangs complementary to the 3’ end of the 50-nt RNA or DNA acceptor oligonucleotides (Fig. 6 and Fig. S8, respectively). In both cases, the reaction occurred at the highest rate and amplitude with a 1-nt 3’ overhang, and the rate decreased with longer 3’ overhangs, despite the complete complementarity of the overhang with the 3’ end of the acceptor. Template switching from the blunt-end starter duplex to the acceptor RNA (0.15 s^−1^) was somewhat slower than from a 1-nt overhang (0.22 s^−1^), but faster than NTA to the same blunt-end duplex at the same dNTP concentration (0.064 s^−1^; Fig. 4B). Similar trends were seen for template-switching to a DNA acceptor, including that template switching from a blunt-end duplex is slower than that from a duplex with a 1-nt overhang but faster than NTA under the same conditions (Fig. S8).

**FIGURE 6.**
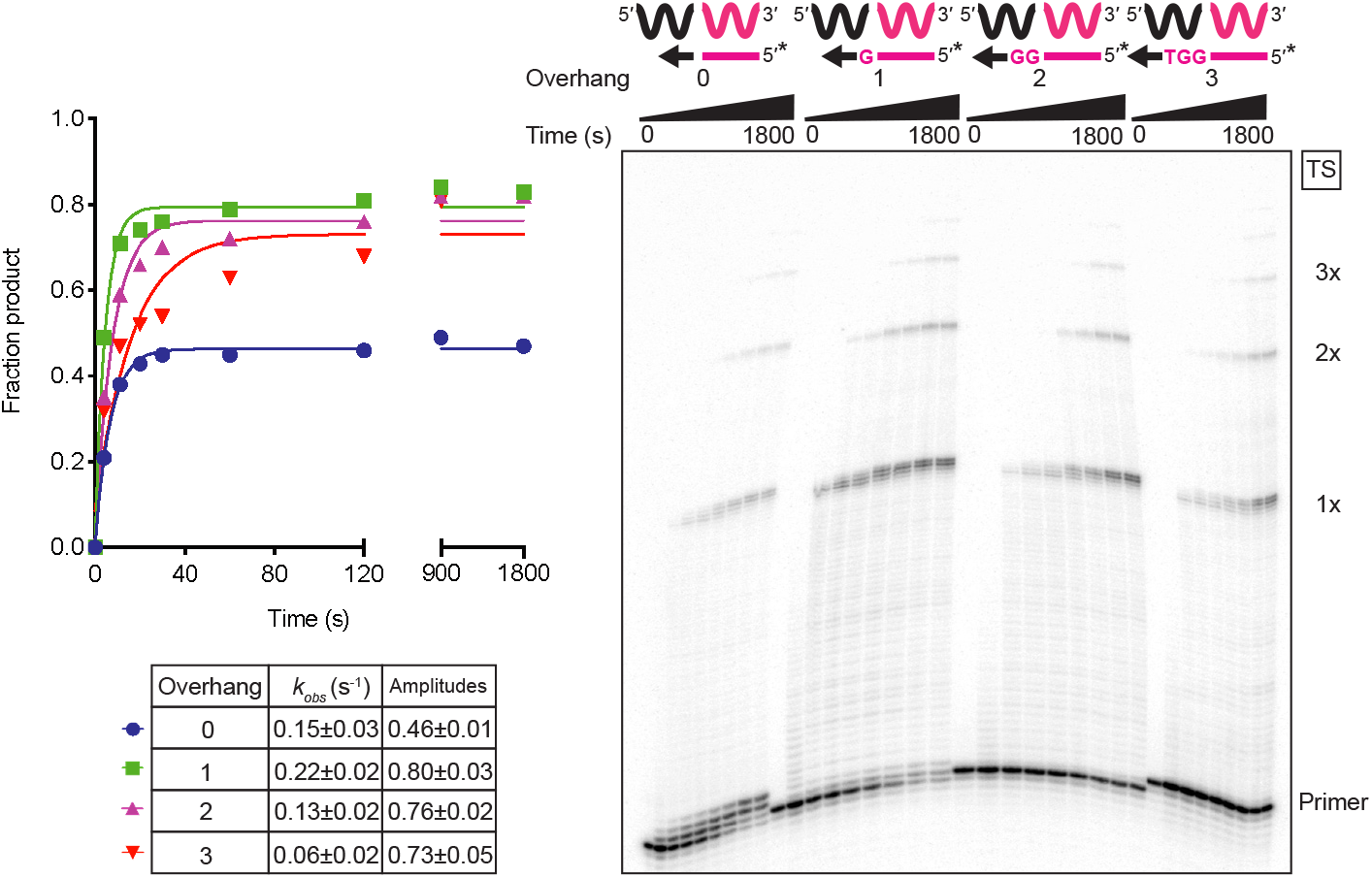
Template-switching is favored by a single nucleotide 3’ overhang. Reactions used 50 nM starter duplexes with a blunt end (0 overhang) or 1, 2, or 3-nt 3’ overhangs, 500 mM GsI-IIC RT, 100 nM 50-nt acceptor RNA in reaction medium containing 200 mM NaCl and were done at 60 °C. Starter duplexes were 5’-^32^P-labeled (*). Reactions were stopped after times ranging from 5 to 1800 s, and the products were analyzed by denaturing PAGE and quantified from phosphorimager scans of the dried gel. The plots show the data fit by a single-exponential function to calculate the *k*_obs_ and amplitude for each time course, and the values are summarized in the table below together with the standard error of the fit. The gel is labeled as in Fig. 1.

Interestingly, the amplitude of template switching from the blunt-end duplex was substantially lower than those for starter duplexes with complementary 1-, 2-, or 3-nt 3’ overhangs, and there was prominent accumulation of products extended by just 1-3 nucleotides, most likely by NTA (see Fig. 6 and Fig. S8 for RNA and DNA acceptors, respectively). The increased NTA observed in the reaction with the blunt-end duplex likely reflects that the absence of a complementary 3’-overhang nucleotide hinders productive binding of the acceptor for template switching, thereby favoring the competing NTA reaction. The lower rates of template switching from starter duplexes with complementary 2- and 3-nt 3’ overhangs could reflect that the template-switching pocket is optimized for the formation of a single base pair that positions the 3’ end of the acceptor for continued cDNA synthesis at the RT active site (see Discussion). NTA products were also lower with these longer 3’ overhangs, as expected for the lower efficiency of NTA to longer 3’ overhangs (see Fig. S5). The finding that the rate of template switching was highest for a complementary 1-nt 3’ overhang indicates that a single base pair is optimal for binding the acceptor in a productive conformation for template switching.

### Template switching fidelity is dictated by a single base pair between the 3’-DNA overhang and the 3’ nucleotide of the acceptor RNA

After determining that a 1-nt 3’ overhang is optimal for template switching, we wanted to compare the kinetics and fidelity of template switching for all four possible base-pair combinations between the 1-nt 3’ overhang and the 3’ nucleotide of the acceptor RNA. As shown in Fig. 7A, template switching occurred efficiently for all four combinations, with indistinguishable rates and slightly higher amplitudes for rC/dG and rG/dC than for rA/dT or rU/dA. The differences in amplitude may be due to a kinetic competition between template switching and the competing NTA reaction, which is sensitive to small differences in the rate of template switching that are not readily detectable in the observed rate constants because of the rapid time scale of the reactions.

**FIGURE 7.**
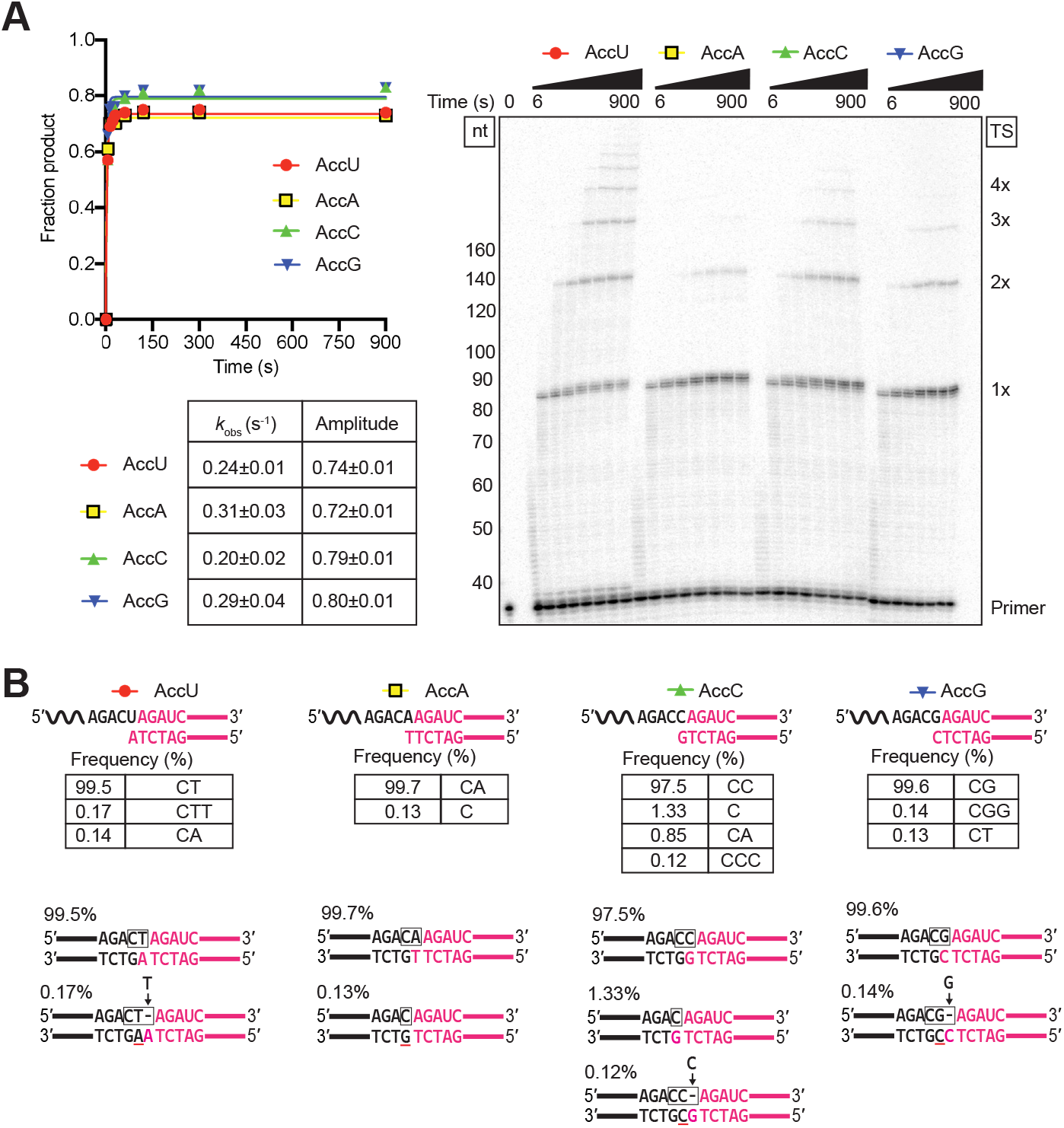
Template switching is directed by a single base pair between the 1-nt 3’ DNA overhang and the 3’ nucleotide of the acceptor RNA. Template-switching reactions with 50-nt acceptor RNAs (100 nM) differing only in their 3’ nucleotide and starter duplexes having a complementary 1-nt 3’ DNA (50 nM) were done with 500 nM GsI-IIC RT at 60 °C in reaction medium containing 200 mM NaCl. *A*, Time courses. Template-switching reactions with ^32^P-labeled starter duplex were stopped after times ranging from 6 to 900 s, and the products were analyzed by denaturing PAGE and quantified from a phosphorimager scan of the dried gel. The gel (right) is labeled as in Fig. 1. The plots (left) show the time course data fit by a single-exponential function to calculate the *k*_obs_ and amplitude for each time course, with the values and standard error of the fit summarized in the table below. *B*, RNA-seq analysis of template-switching junctions. Template-switching reactions with unlabeled starter duplexes were done under the same conditions as in panel A with the reaction stopped after 15 min. RNA-seq analysis was done as described in Fig. 1 and Experimental Procedures. The figure diagrams the template-switching reaction for each acceptor/starter duplex combination above, with the sequences and percentages of the most frequent template-switching junctions (≥0.1%) for each combination listed below. Nucleotides derived from the acceptor are in black, and nucleotides derived from the starter duplex are in red, with the box indicating the junction sequence. A black letter with red underline indicates a nucleotide inferred to result from NTA to the 3’ end of the DNA primer. A gap in the top strand due to NTA is shown as a dash, and nucleotides inferred to fill the gap after PCR to add RNA-seq adapters (Fig. 1A) are shown above the line with an arrow pointing to the gap. The low frequency junctions containing an extra nucleotide (CTT for the 3’ U acceptor (0.17%), CCC for the C acceptor (0.12%), and CGG for the 3’ G acceptor (0.14%)) can be explained by template switching from donors that have undergone an NTA of a complementary nucleotide resulting in a 2’-nt 3’ overhang that leaves a gap in the top strand, which is filled by a complementary nucleotide during the PCR used to add RNA-seq adapters. Other aberrant products may reflect heterogeneity or resections at the 3’ ends of the synthesized oligonucleotides (*e.g.*, the 3’ C junction for the 3’ C acceptor (1.33%) can be explained by template switching to a mis-synthesized or resected acceptor RNA lacking the terminal C residue. Complete data for junction sequences are shown in Table S3.

Sequencing of the template-switching junctions between the starter duplex and the acceptor RNA showed accurate extension of all four combinations, with 97.5 to 99.7% of the products having the seamless junctions expected for extension of the single base pair (Fig. 7B). The sequencing also showed a smattering of aberrant products due to the competing NTA reaction or impurities in the synthetic oligonucleotides, as described in the legend of Fig. 7B. The high efficiency of seamless template switching dictated by a single base pair between the 1-nt 3’ overhang and the 3’ nucleotide of the acceptor presumably reflects a very tight and specific interaction of complementary 3’ termini within the binding pocket for the acceptor RNA.

### Nucleotide sequence biases in TGIRT-seq reflect the efficiency of template switching to different 3’ nucleotides of acceptor RNAs at high salt concentration

Previous TGIRT-seq analysis of miRNA reference sets showed that 3’-sequence biases in TGIRT-seq are almost completely confined to the 3’-terminal nucleotide of the acceptor, with a marked preference for miRNAs with a 3’ G residue and against those with a 3’ U residue (49). In light of our finding that template switching is slower at high salt concentrations (Fig. 2), we wondered whether the high salt concentration used in TGIRT-seq to suppress multiple template switches (450 mM NaCl (8, 9)) might exacerbate differences between acceptor RNAs with different 3’ nucleotides, which are muted at 200 mM NaCl (Fig. 7).

To test this idea, we compared template switching to acceptor RNAs with all four Watson-Crick base-pairing combinations between the single-nucleotide 3’ overhang and 3’ nucleotide of the acceptor at 450 mM NaCl. Under these conditions, we found larger differences in the amplitude of the reaction for acceptors with different 3’ nucleotides in order G>C≈A>U (Fig. 8A), which match the previously determined nucleotide sequence biases for TGIRT-seq (49). Additionally, when the reactions were carried out with lower concentrations of acceptor RNAs (10 nM) to more closely resemble the conditions during TGIRT-seq, the difference in amplitude for template switching to the most and least favored 3’ nucleotides (G and U, respectively) increased (Fig. 8B), as expected if productive binding of the acceptor became limiting under these conditions. The more uniform rates and amplitudes for template-switching to acceptor RNAs with different 3’ nucleotides at 200 mM NaCl (Fig. 7) suggests that the nucleotide sequence biases for 3’ adaptor in TGIRT-seq might be ameliorated by carrying out the initial template-switching step at a lower salt concentration.

**FIGURE 8.**
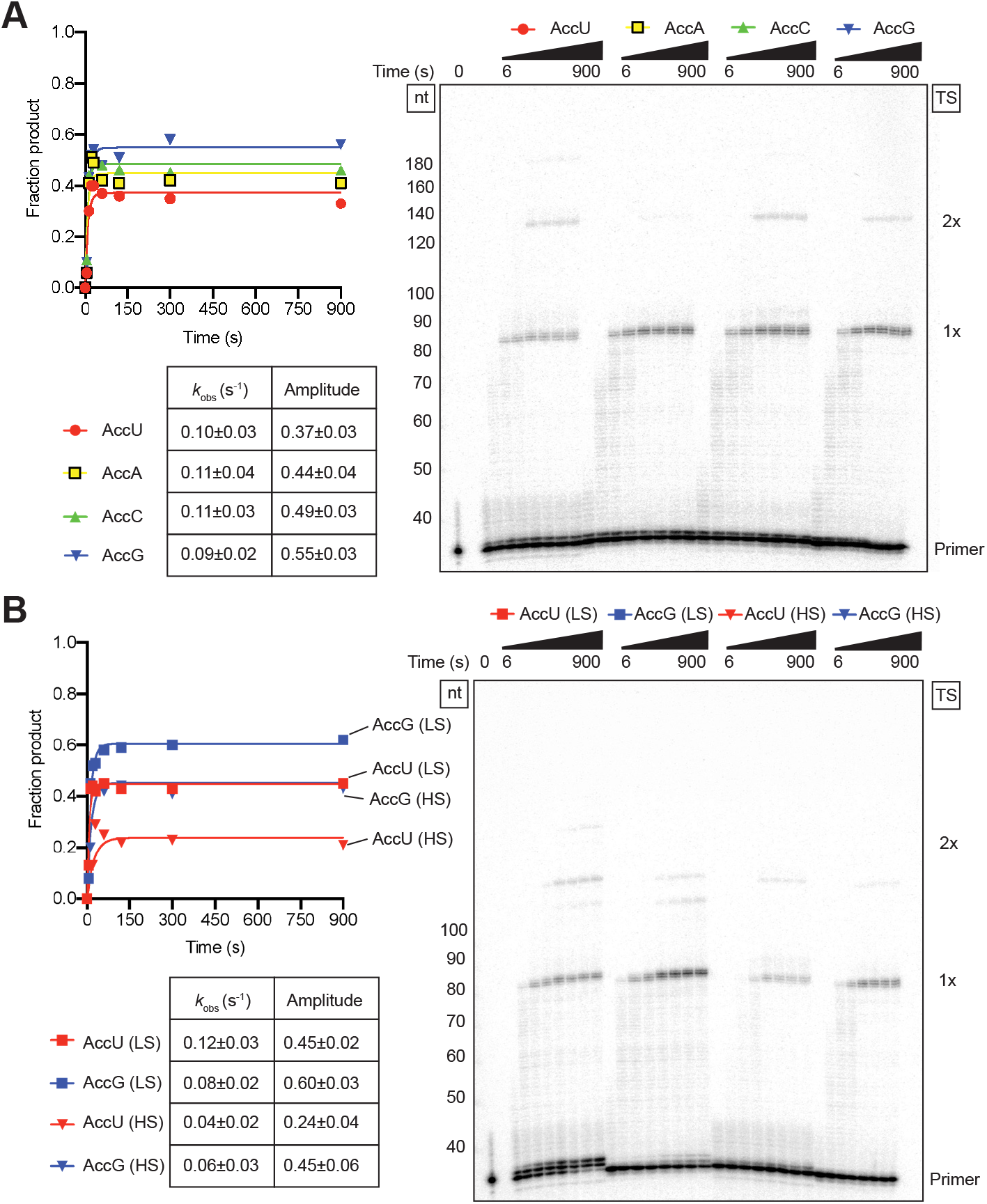
Biases in 3’-adapter addition in TGIRT-seq reflect the efficiency of template-switching to acceptor RNAs with different 3’ nucleotides at a high salt concentration. *A*, Template-switching reactions with 50-nt acceptor RNAs differing only in their 3’ nucleotide and ^32^P-labeled starter duplexes having a complementary 1-nt 3’ DNA overhang were done as in Fig. 7, but in reaction medium containing 450 mM instead of 200 mM NaCl. *B*, Template-switching reactions were done as in panel A, but with 10 nM instead of 100 nM acceptor RNA in reaction medium containing either 200 mM NaCl (LS) or 450 mM NaCl (HS). The gels are labeled as in Fig. 1. The plots to the left of the gel show the time course data fit by a single-exponential function to calculate the *k*_obs_ and amplitude for each time course, and the values and standard errors of the fit are summarized in the tables below the plots.

### Template switching from a 1-nt overhang duplex disfavors the extension of mismatches

To further investigate how GsI-IIC RT template-switching discriminates between complementary and mismatched acceptors, we carried out parallel reactions comparing a complementary 3’-C acceptor and 3’-G overhang combination to a mismatched 3’-C acceptor and 3’-C overhang combination (Fig. 9). Both the rate and amplitude of product formation were substantially decreased for the mismatched combination, with the rate ~8-fold lower (*k*_obs_ = 0.03 compared to 0.23 s^−1^) and the amplitude ~50% lower than those for the complementary combination (0.43 compared to 0.88, although the slow phase appeared to be continuing past 900 s for the mismatched combination; Fig. 9A).

**FIGURE 9.**
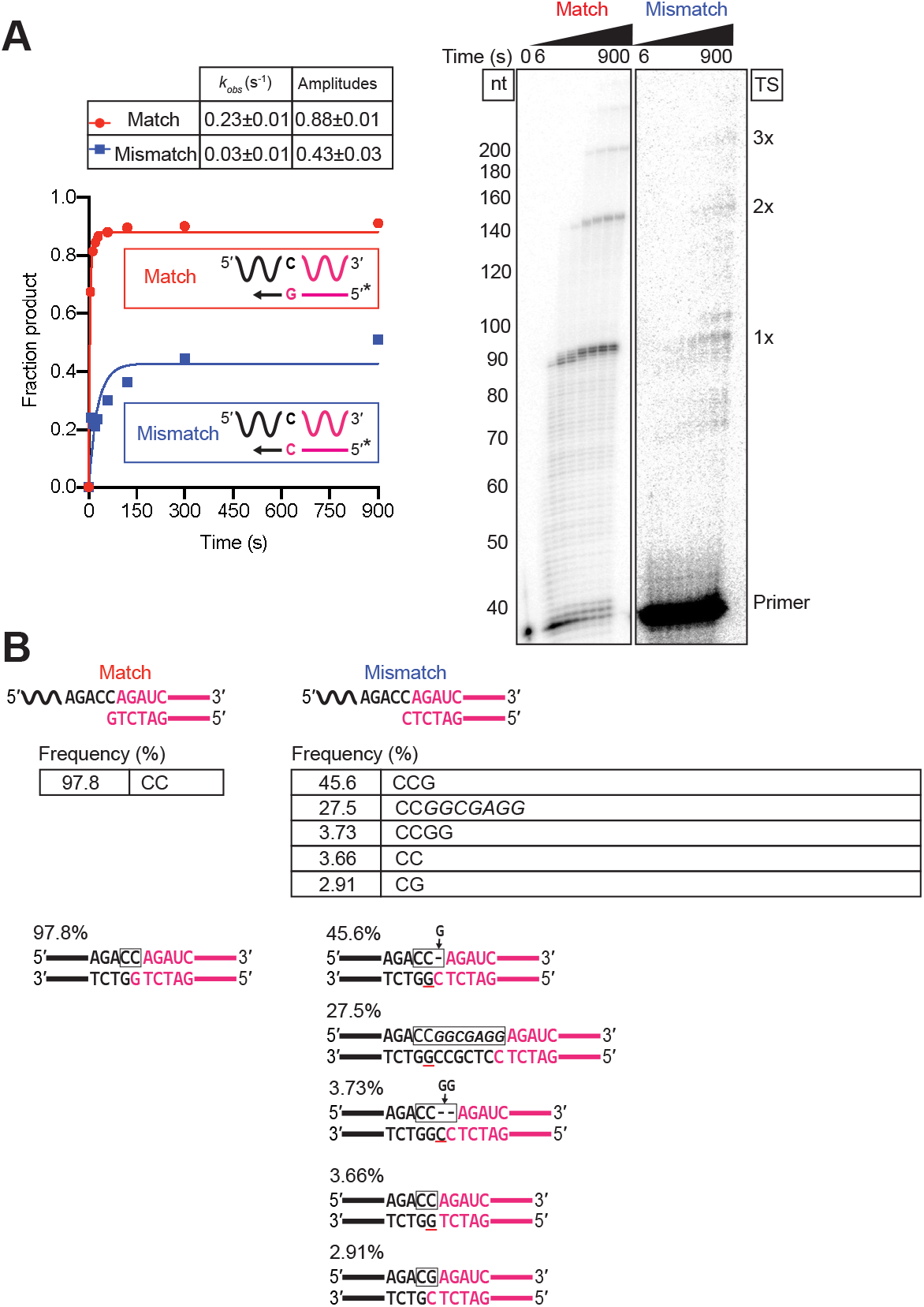
Template switching by GsI-IIC RT disfavors the extension of mismatches. Templateswitching reactions with starter duplex have a 1-nt 3’ C overhang and otherwise identical 50-nt acceptor RNAs having a complementary (matched) 3’ G or non-complementary (mismatched) 3’ C were done as described in Fig. 7. *A*, Time courses. Template-switching reactions with ^32^P-labeled starter duplex were stopped after times ranging from 6 to 900 s, and the products were analyzed by denaturing PAGE and quantified from phosphorimager scans of the dried gel. The gels (different exposures for the matched and mismatched configurations) are labeled as in Fig. 1. The plots to the left of the gel show the data fit by a single-exponential function to calculate the *k*_obs_ and amplitude for each time course, and the values together with the standard error of the fit are summarized in the table above. *B*, RNA-seq analysis of template-switching junctions. Template-switching reactions with unlabeled starter duplexes were done under the same conditions as in panel A with the reaction stopped after 15 min. RNA-seq analysis was done as described in Fig. 1 and Experimental Procedures. The figure diagrams the template-switching reaction for each acceptor/starter duplex combination above, with the sequences and percentages of the most frequent template-switching junctions for the mismatched combination shown below. Nucleotides that were derived from the acceptor are in black, and nucleotides derived from the starter duplex are in red, with the box indicating the junction sequence. A black letter with red underline indicates a nucleotide residue inferred to result from NTA. A gap in the top strand due to NTA is shown as a dash, and nucleotides inferred to fill the gap after PCR to add RNA-seq adapters are indicated above the line with arrows pointing to the gap. Nucleotides in italics are putatively derived from an intermediate template switch to contaminating oligonucleotides present in the enzyme preparation or reagents. Complete data for junction sequences are shown in Table S3.

Sequencing of the reaction products again showed high fidelity for the complementary combination, with 97.8% of the product having the sequence expected for base pairing of the 3’-G overhang nucleotide to the 3’ C of the acceptor (Fig. 9B). By contrast, the most abundant template-switching junction for the mismatched combination was CCG (45.6%), which can be explained by template switching from a donor that has undergone NTA to add a complementary G overhang, leaving a gap (dash) in the template strand, which is filled by a complementary nucleotide (indicated by an arrow pointing to the gap) during the PCR used to add RNA-seq adapters (Fig. 9B). Surprisingly, the second most abundant junction (27.5%) contained a relatively long insertion that appears to reflect an intermediate template switch to a low-level contaminating oligonucleotide (italics) with a 3’ G that could base pair to the 3’-C overhang of the starter duplex. Another aberrant junction (CCGG; 3.73%) required two NTAs before adding a complementary G overhang, and another (CC, 3.66%) can be explained by NTA of a complementary G overhang to a bunt-end version of the starter duplex that may be present at low levels due to mis-synthesis or resection of the mismatched 3’ overhang. By contrast, the junction expected for extension of the mismatch (CG; with the G in the top strand incorporated opposite the mismatched C in the bottom strand after PCR) was detected at lower frequency (2.91%). The preference for these inefficient alternative reactions, which revert to a complementary 3’-overhang nucleotide for template switching rather than use a mismatched acceptor RNA added in excess, indicates that GsI-IIC has a strong aversion to extending mismatches at the template-switching junction.

### Mechanisms of template switching from a blunt-end duplex

Finally, to explore the mechanism of template switching from a blunt-end duplex, we performed similar template-switching reactions from the blunt-end version of the starter duplex to acceptor RNAs ending in U, A, C, G, and N, where N is an equimolar mixture of the four acceptor RNAs.

As shown in Fig. 10A, the rate and amplitude of template switching starting from an artificial blunt-end duplex were highest for the 3’-N acceptor, which can base pair with any 3’-overhang nucleotide added by NTA, and for the 3’-U acceptor, which can base pair with a 3’-overhang A, the preferred nucleotide for NTA to the same blunt-end duplex (see Fig. 5 and Fig. S5 for NTA preferences). The efficiency of template switching to the remaining acceptors followed the order C>G>A, roughly matching the order expected from the efficiencies of NTA to the blunt-end duplex (G>C≈T; Fig. 5A). The efficient use of the 3’ U acceptor for template switching from a blunt-end duplex contrasts with its least efficient use for template-switching from a complementary 1-nt 3’ overhang duplex (Figs. 7 and 8) and indicates that NTA of a complementary A overhang that can base pair with the 3’ U is rate limiting for template switching from the blunt-end duplex.

**FIGURE 10.**
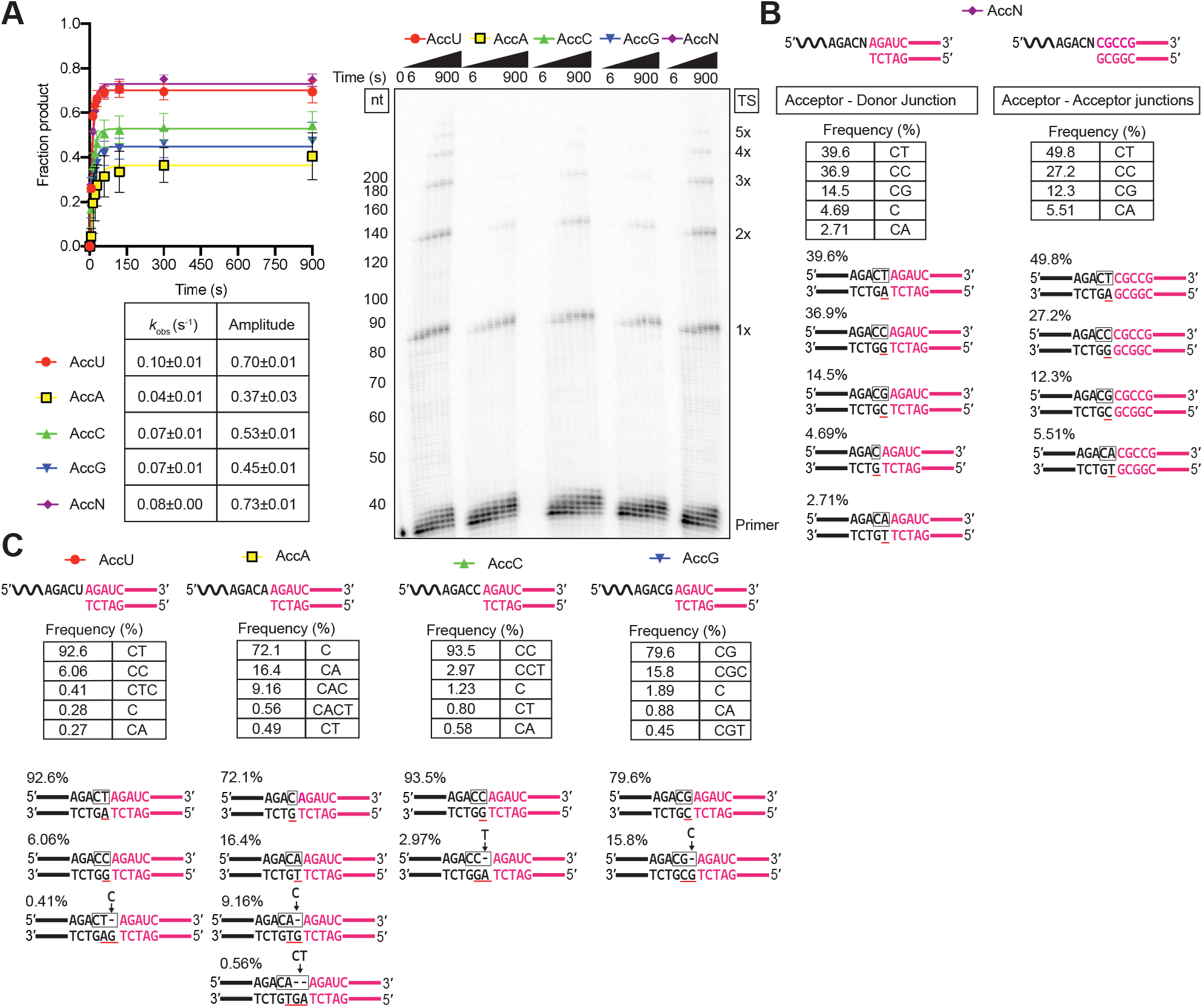
Template switching from a blunt-end starter duplex is inefficient and yields heterologous junction sequences. Template-switching reactions with 50-nt acceptor RNAs (100 nM) differing only in their 3’-nucleotide residue and a blunt end-starter R2 RNA/R2R DNA starter duplex (50 nM) were done as described in Fig. 7. The 3’ N acceptor is an equimolar mix of acceptor RNAs having 3’ A, C, G, or U residues added at the same total concentration (100 nM) as each individual acceptor. *A*, Time courses. Template-switching reactions with ^32^P-labeled starter duplex were stopped after times ranging from 6 to 900 s, and the products were analyzed by denaturing PAGE and quantified from phosphorimager scans of the dried gel. The gel is labeled as in Fig. 1. The plot to the left of the gel shows the data for each acceptor fit by a single-exponential function, with the values and standard error of the fit summarized in the table below the plot. *B* and *C*, RNA-seq analysis of template-switching junctions between the acceptor RNAs and unlabeled blunt-end duplex were done for 15 min under the same conditions as in panel A. The figure diagrams the template-switching reaction for each combination, with the sequences and frequencies of the most frequent template-switching junctions shown below. Panel B shows sequences and frequencies for both the initial Acceptor-Donor junctions and subsequent Acceptor-Acceptor junctions, while panel C shows only those for the initial Acceptor-Donor junctions. Nucleotides that were derived from the acceptor RNA are in black, and nucleotides derived from the synthetic blunt-end duplex or blunt-end duplexes formed after completion of cDNA synthesis are in red, with the box indicating the junction sequence. A black letter with red underline indicates a nucleotide inferred to result from NTA to a blunt-end duplex. Gaps in the top strand due to NTA are shown as dashes, and nucleotides inferred to fill those gaps after PCR to add RNA-seq adapters are indicated above the line with arrows pointing to the gap. Complete data for junction sequences are shown in Table S3.

Additional experiments showed that template switching to a 3’-N acceptor had a higher *K*_1/2_ for dNTP from a blunt-end duplex than a from 1-nt overhang 3’ N duplex (0.92 mM and 0.03 mM, respectively; Fig. S9A, and Fig. S10 (A and B)), and similar results were obtained for template switching to a 3’-C acceptor from a blunt-end or 3’-G-overhang duplex with varying concentrations of dGTP, the nucleotide complementary to the first and second base in the acceptor RNA (*K*_1/2_ = 3 and 0.03 mM, respectively; Fig. S9B, and Fig. S10 (C and D)). This >30-fold higher *K*_1/2_ for dNTP addition for template switching from a blunt-end duplex is similar to that for NTA (Figs. 4 and 5). While this finding is consistent with a requirement for NTA to produce a complementary 1-nt 3’ overhang prior to template switching, it could also reflect template switching directly from a blunt-end duplex, with weaker dNTP binding reflecting that the templating 3’ nucleotide of the acceptor is not anchored by a phosphodiester bond to the downstream template strand as it would ordinarily be for cDNA synthesis.

To attempt to distinguish these possibilities, we analyzed the junctions resulting from template switching from the blunt-end duplex to the 3’-N acceptor by RNA-seq (Fig. 10B). For this analysis, we separately analyzed the two types of template-switching junctions formed in the reaction: those resulting from template switching from the starter duplex to the 3’ end of the acceptor RNA (Acceptor-Donor Junctions) and those resulting from secondary template switches from the 5’ end of an acceptor RNA after completion of cDNA synthesis to the 3’ end of another acceptor RNA Acceptor-Acceptor Junctions). The expectation was that template-switching directly from a blunt-end duplex would results in a roughly equal proportions of junctions for 3’ A, C, G, and U acceptors, while template switching via NTA would parallel the efficiency of NTA of the complementary 3’-overhang nucleotide to the blunt end duplex. For both Acceptor-Donor and Acceptor-Acceptor junctions, the proportion of junctions for different acceptor 3’-nucleotides was U>C>>G>A (Fig. 10B, right), roughly paralleling that for NTA of the complementary nucleotide (A>G>>C≈T) to the same blunt end-duplex (Fig. 5A). The proportion of CT junctions, requiring NTA of a 3’ overhang A that can base pair to the 3’ U acceptor was somewhat higher for the secondary template switches from the 5’ end of the acceptor than those from the artificial starter duplex. Additionally, template switching from the blunt-end starter duplex yielded a significant proportion of single C junctions (4.69%), possibly via NTA of a G residue to a population of acceptors ending with a single 3’ C instead of a 3’ CN (also seen in the experiments below). Together, these findings suggest that template switching from a blunt-end duplex occurs largely via NTA to produce a 3’-overhang nucleotide that can base pair with the 3’ nucleotide of the acceptor.

Additional support for this conclusion came from template-switching reactions from a blunt-end duplex to individual acceptor RNAs with A, C, G or U 3’ nucleotides (Fig. 10C). In this case, the expectation for the NTA-mediated mechanism was that that the highest proportion of seamless junctions would be found for the 3’ U and C acceptors, which require NTA of a complementary A or G overhang, the two residue most favored for NTA to the blunt-end duplex. By contrast, the lowest proportion should be found for the 3’ G and A acceptor, which require NTA of a complementary or C or U overhang, the two residues least favored for NTA to the blunt-end duplex. In agreement with these expectations, the 3’ U and C acceptors yielding seamless CT or CG junctions at frequencies of 92.6% and 93.5%, respectively, and the 3’ G and 3’ A acceptors yielding seamless CG and CA junctions at frequencies of 79.6 and 16.4%, respectively (Fig. 9C). The CC junction seen at 6.06% for the 3’-U acceptor could also reflect NTA of a G overhang, enabling a G:U wobble base pair with the 3’ U (50), with the subsequent PCR inserting a C opposite the G added by NTA.

All four of the acceptors yielded some junctions in which one or two NTAs occurred prior to the template switch. Additionally, all four acceptors yielded some junctions with a single C residue, suggesting either that acceptor RNAs lacking the 3’ terminal nucleotide are present at low concentrations in all four acceptor RNA preparations or that some mechanism exists for resection of the 3’-terminal residue of the acceptor. In the case of the 3’-A acceptor, the single C junction was the most abundant (72.1%), possibly reflecting that NTA of a G overhang complementary to the 3’ C of this rare acceptor is much more efficient than NTA of a 3’ U overhang complementary to the 3’ A of the predominant acceptor.

Considered together, our results indicate that template switching from a blunt-end duplex occurs largely via NTA to add a complementary 3’ overhang nucleotide to the starter duplex. However, our results do not exclude the possibility that some template switching can occur directly from a blunt-end duplex without NTA, particularly in those cases for which NTA of a complementary 3’-overhang nucleotide to the blunt-end starter duplex is inefficient (see Discussion).

## Discussion

Here, we analyzed the end-to-end template switching and NTA activities of the thermostable GsI-IIC group II intron RT. Our results indicate that template switching by this enzyme is favored by a single base pair between the 3’ nucleotide of the acceptor nucleic acid and a 1-nt 3’-DNA overhang of an RNA/DNA heteroduplex, with the 1-nt overhang provided either as part of an artificial starter duplex or by NTA to the 3’ end of a cDNA after the completion of cDNA synthesis. Mechanistically, there are both shared features and differences with the end-to-end template switching activities of other non-LTR-retroelement RTs, as discussed further below. The finding that a single base pair is sufficient to faithfully direct template switching at 60 °C, the operational temperature for GsI-IIC RT, indicates a very high degree of specificity, and presumably affinity, for this base pair within the template-switching pocket, and it has implications for both the biological function and biotechnological applications of template switching (see below).

### Template switching and non-templated nucleotide addition are related reactions

Fig. 11A shows a model of the template-switching reaction after cDNA synthesis from a template RNA with a 5’ A residue. After addition of a deoxythymidine to complete cDNA synthesis, the enzyme releases pyrophosphate and undergoes translocation (51), moving the terminal rA/dT base pair out of the RT active site (*i.e.*, from position −1 to +1). This translocation leaves the GsI-IIC RT active site (position −1) free to bind and add a nucleotide by NTA. Any of the four deoxynucleotides can be added by NTA, but our data reveal a preference for dA. Upon incorporation of a nucleotide and pyrophosphate release, translocation of the added A residue from −1 to +1 creates a binding pocket that favors base pairing of an acceptor RNA with a complementary 3’ U residue, resulting in positioning of the penultimate C residue of the acceptor as the templating RNA base at position −1 for continued cDNA synthesis.

**FIGURE 11.**
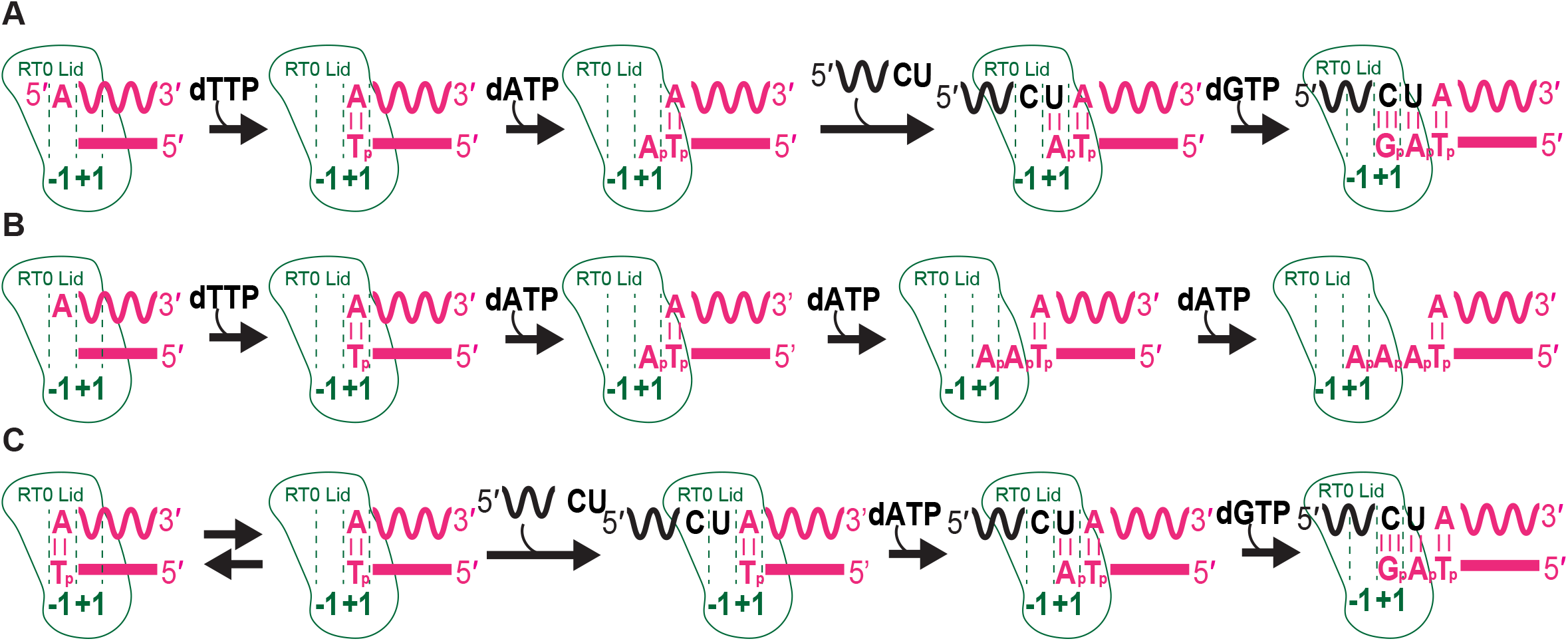
Models of template switching and non-templated nucleotide addition reactions. *A*, Template-switching to an acceptor RNA from an RNA template/DNA primer heteroduplex with a 1-nt 3’ overhang added by NTA after completion of cDNA synthesis or as an artificial starter duplex. *B*, NTA to a blunt-end RNA/DNA heteroduplex in the absence of an acceptor nucleic acid. *C*, Template-switching to an acceptor RNA from a blunt-end RNA/DNA heteroduplex without NTA. See Discussion for details.

A similar mechanism likely applies for artificial starter duplexes with a 1-nt 3’ overhang. In this case, it is unclear whether the 3’-overhang nucleotide is initially bound at position −1 (the RT active site) or position +1 (the position occupied by a 1-nt 3’ overhang after NTA; see Fig. 11). As above, a 1-nt 3’-overhang that is initially in the +1 position would create a binding pocket for an acceptor RNA with a complementary 3’ nucleotide, leaving the penultimate 3’ residue of the acceptor in the −1 position to serve as the templating base for cDNA synthesis. Alternatively, a 1-nt 3’ overhang that is initially bound in the −1 position would have to translocate to the +1 position before polymerization. Mechanistically, the two positions of the 3’-overhang may be in an equilibrium that is shifted by base pairing of the acceptor and the continuation of cDNA synthesis, a feature that has been described for Y-family DNA polymerases (discussed further below) (47, 52, 53). A comparison of artificial starter duplexes with 1-nt 3’ overhangs that can or cannot base pair to the 3’ nucleotide of the acceptor indicates that binding of a mismatched acceptor at the +1 position and/or extension from a mismatch at the −1 position are strongly disfavored (Fig. 9).

In the absence of an acceptor template with complementarity to the 1-nt 3’ overhang of the primer, the RT can add additional nucleotides by NTA (Fig. 11B). The second and third NTA reactions are progressively slower than the first (Fig. 4 and Fig. 5), increasing the time window for sampling potential acceptor templates for complementarity. Longer base-pairing interactions assist template switching by retroviral and LTR-retrotransposon RTs (18), but the opposite is true for GsI-IIC RT, with overhang lengths beyond 1-nt decreasing the efficiency of template switching (Fig. 6 and Fig. S8). This difference may reflect that template-switching by retroviral RTs involves partial or complete dissociation of the enzyme from the initial template, enabling the completed cDNA to base pair to the new template for reinitiation of cDNA synthesis by the same or another RT. By contrast, non-LTR-retroelement RTs bind the donor and acceptor simultaneously without dissociating, leaving a relatively small binding pocket that may sterically hinder access to multiple overhang bases (41).

### Non-templated nucleotide addition in the absence of an acceptor nucleic acid

In the absence of an acceptor with a complementary 3’ nucleotide, NTA continues for up to three (and occasionally four) nucleotides, progressively translocating the completed cDNA/RNA template duplex away from the active site (Fig. 11B). Because group II intron RTs lack RNase H activity and bind RNA/DNA heteroduplexes tightly, such translocation via NTA may facilitate dissociation of the enzyme from the completed cDNA product when an acceptor nucleic acid for template switching is unavailable (34).

Our biochemical analysis shows that in the absence of an acceptor nucleic acid, NTA by GsI-IIC RT favors purine residues, particularly an A residue, at all steps (Fig. 5). A similar nucleotide preference, termed the ‘A-rule’, has been found for many other polymerases, albeit with some notable exceptions (16, 43, 54) and is thought to reflect more stable base-stacking of purines on the terminal base pair of the completed duplex (55). The higher *K*_1/2_ of dNTP addition for NTA compared to reverse transcription suggests a low affinity interaction when compared to the extension reaction (compare Fig. 4 and Fig. S10, A and B, right panels), presumably due to the absence of base pairing with a templating base. The pausing step we elucidated by modeling of processive NTA addition could indicate (i) the presence of a pre-insertion step, during which the blunt-end duplex occupies the −1 nucleotide-binding site, effectively inhibiting nucleotides from binding; (ii) backtracking of the RT, a process documented for retroviral RTs (56, 57); or (iii) enzyme dissociation from the template-primer heteroduplex. Structural and kinetic analysis of Y-family DNA polymerases showed that an analogous pausing step reflects a dynamic equilibrium between the 3’ nucleotide of primer occupying the −1 position in the pre-insertion state and the nucleotide-binding competent +1 position (47, 52, 53). It is possible that NTA preferences for different nucleotides may be influenced by the sequence of the starter duplex, as a similar trend but with a stronger preference for A over G residues was found for NTA to the 3’ end of completed cDNAs in TGIRT-seq of miRNA reference sets containing 962 human miRNAs (A, 86.6%; G, 9.9%; C, 2.0%; and U, 1.4% for the first NTA, with 83.3% of the NTAs being a single nucleotide (SRA accession number NTT1, SRX5005854).

### Template switching from a blunt-end duplex

Our analysis of junction sequences suggests that template switching of GsI-IIC RT from a blunt-end duplex occurs largely via NTA of a 3’-overhang nucleotide that can base pair to the 3’ nucleotide of the acceptor (Fig. 10). However, the finding that the rate of template switching from a bunt-end duplex to both RNA and DNA acceptor is faster than the rate of NTA to same duplex (Fig. 6 and Fig. S8) could reflect that some template-switching can also occur directly from a blunt end duplex without NTA.

As shown in Fig. 11C, template switching directly from a blunt duplex would require that the terminal base pair of the duplex be located at the +1 position, thereby allowing binding of the terminal nucleotide of the acceptor at the RT active site (position −1). The latter then forms a binding pocket for a complementary dNTP, which can be polymerized onto the DNA primer in a normal manner (Fig. 11C). Unlike the situation after the completion of cDNA synthesis (Fig. 11A), the binding or movement of the terminal base pair of an artificial blunt end duplex to the +1 position is not linked to NTA) and could again reflect a natural equilibrium between the −1 and +1 position, which is pushed forward by binding of the acceptor and the continuation of cDNA synthesis.

We note that the junction sequences formed by direct template switching from a blunt-end duplex would be indistinguishable from those formed via NTA of a nucleotide complementary to the 3’ end of the acceptor, the only difference being whether the 3’ nucleotide of the acceptor served as the templating base. The finding that the *K*_1/2_ for dNTP for template-switching from a blunt-end duplex is >30-fold higher than that for template switching from a 1-nt 3’ overhang duplex (Fig. S10) could reflect either a requirement for NTA or weaker dNTP binding opposite the 3’ nucleotide of the acceptor RNA that is not fixed in position by a base pair to a 1-nt overhang or by a phosphodiester bond to the downstream RNA template strand as it would ordinarily be during cDNA synthesis.

### Comparison of GsI-IIC RT to other non-LTR-retroelement RTs

Non-LTR-retroelement RTs typically have an NTE with a conserved RT0 motif that likely contributes to a structurally similar binding pocket for the 3’ end of the acceptor nucleic acid during template switching (5, 19, 41, 42). Thus, it is unsurprising that many features of the template switching and NTA mechanisms of Gs-IIC RT are similar to those found previously for the MRP and R2 element RTs, including a prominent role for NTA in adding nucleotides to the 3’ end of the cDNA that can base pair to the 3’ end of acceptor RNAs (25, 33, 34). For the R2 RT, biochemical experiments examining NTA after the completion of cDNA synthesis found similar nucleotide preference A>G>>C≈T, with NTA limited to 1-4 nt and with longer 3’ overhangs disfavoring template switching (33, 34), all similar to our findings for GsI-IIC RT.

Despite these similarities, there are significant differences in template switching by the three RTs that are relevant to biotechnological applications and suggest how template-switching activity may be adapted to fit the life cycle of different non-LTR-retroelements. One important difference is that the R2 RT appears to require a ‘running start’ to template switch and is unable to rebind to a completed cDNA with a 3’ overhang added by NTA (33, 34). In contrast, GsI-IIC RT can bind to and template-switch efficiently from synthetic RNA template/DNA primer duplexes with a 1-nt 3’ overhang, an ability that enables precise RNA-seq adapter addition in TGIRT-seq (6). Further, unlike GsI-IIC RT, which requires an RNA/DNA heteroduplex with a 3’ overhang for efficient template-switching, the R2 element RT can initiate reverse transcription near the 3’ end of an acceptor RNA by using an RNA primer (33), and the MRP RT can do so using a singlestranded DNA primer (25). Finally, unlike GsI-IIC RT template switching, the specificity of which is dictated by a single base pair to the 3’ nucleotide of the acceptor, the MRP RT has a strong preference for template switching to RNAs with a 3’ CCA and to initiate cDNA synthesis opposite the penultimate C residue (C-2), even in the absence of complementarity to the DNA primer (25). These features likely reflect adaptation of the MRP RT template-switching activity to initiate at the 3’ tRNA-like structure of the plasmid transcript for plasmid replication.

### Possible biological functions of non-LTR-retroelement RT template switching

The high efficiency of end-to-end template switching by non-LTR-retroelement RTs and its dependence upon a conserved binding pocket that is not present in retroviral RTs suggest that this activity has an important biological function. In the case of insect R2 and mammalian LINE elements, template switching has been proposed to function in the attachment of the 3’ end of a cDNA that was initiated by TPRT to upstream host DNA sequences (35, 37). While this remains an attractive hypothesis, other modes of DNA attachment have not been excluded, and experiments investigating the retrohoming of a linear group II intron RNA in *Drosophila melanogaster*, an analogous situation requiring attachment of a free 3’ cDNA to upstream host sequences, indicated involvement of non-homologous end-joining enzymes rather than template switching (39). Additionally, recent findings indicate that chimeric integration products of human LINE-1 elements can result from RNA ligation rather than template switching (58).

A different and not mutually exclusive hypothesis is suggested by the common requirement of all non-LTR-retroelement RTs to synthesize full-length cDNAs of long RNA templates that are vulnerable to oxidative damage and nicking by host endonucleases (*e.g.*, RNase E in the case of bacterial group II intron RTs; (59)). Template switching may provide a means of reverse transcribing through such a nicked RNA template *in vivo*, enabling the synthesis of long cDNAs from discontinuous RNA templates, and would disfavor multiple NTAs in functionally important sequences, accounting for the kinetic preference for a single NTA. We note that template switching may play a similar role for RNA viruses, whose RNA-dependent RNA polymerases (RdRPs) are closely related to group II intron RTs, including a structurally homologous acceptor RNA binding pocket, termed motif G in viral RdRPs (41, 60). This central function may be combined with additional functions specific to the life cycle of different non-LTR-retroelements and viruses, such as preferential template-switching to a 3’-tRNA-like structure in the cases of MRP RTs (25) and higher rates of internal template switches to generate recombinant viruses to evade host defenses or repair defective genomes (61).

### Implications for RNA-seq

Finally, our results have significant implications for the use of group II intron RT template switching in RNA-seq. First, we found that the configuration of starter RNA template/DNA primer duplex used in TGIRT-seq in which a 1-nt 3’ DNA overhang base pairs to the 3’ nucleotide of target RNAs (Fig. 1A), is in fact the most accurate and efficient configuration for GsI-IIC RT template switching. The thermostable TeI4c group II intron RT has also been shown to template-switch efficiently from an artificial starter duplex with a 1-nt 3’ overhang, and this is likely to be a common characteristic of group II intron RTs (6). Second, we found that the previously determined 3’-sequence biases in TGIRT-seq reflect the efficiency of the template switching to acceptor RNAs with different 3’ nucleotides, particularly under the high-salt conditions used for TGIRT-seq (Fig. 8). These 3’ biases, which are restricted to 3’ terminal nucleotide of the acceptor, account for about half the sequence bias in TGIRT-seq, the remainder coming from the 5’ App RNA/DNA ligase used for R1R adapter ligation (see Fig. 1A)(49). Although we showed previously that these 3’ biases could be remediated computationally or by changing the ratio of 3’ DNA overhangs in the starter duplex (49), we found here that they can be more simply moderated by carrying out the template-switching reaction at lower salt concentrations (200 mM NaCl; Fig. 8). It should also be noted that these 3’-end biases for template-switching by GsI-IIC RT are not universal, as TeI4c RT preferentially template switches to acceptor RNAs with a 3’-A residue (Yidan Qin and A.M.L., unpublished data), implying that these template-switching biases can be modified and perhaps eliminated by protein engineering of the acceptor-binding pocket. Finally, in addition to reducing template-switching biases, we found that lower salt concentrations substantially increase the efficiency of TGIRT template switching (2-4-fold increase in rate; 1.7-fold increase in amplitude; Fig. 2), perhaps by stabilizing a productive enzyme-substrate conformation. The higher efficiency of template switching at lower salt concentrations enables more efficient capture of RNA templates (Fig. S3), as well as more efficient use of RNAs with modified 3’ ends, including 3’ phosphate and 2’-*O*-Me groups (Fig. 3), which are present at the 3’ ends of piRNAs and plant miRNAs (62, 63). Although these benefits for different applications must be weighed against the effect of increased multiple template switching at lower salt concentrations, recent TGIRT-seq of human plasma RNA at 200 mM instead of 450 mM NaCl substantially increased the efficiency of TGIRT-seq library construction, without unacceptably increasing the proportion of multiple template-switches (≤2.5% fusion reads, which include multiple-template switches; Wu et al., manuscript in preparation).

## Experimental Procedures

### Protein purification

GsI-IIC RT with an N-terminal maltose-binding protein tag to keep the protein soluble in the absence of bound nucleic acids was expressed from plasmid pMRF-GsI-IIC and purified with minor modifications of a previously described procedure (6). A freshly transformed colony of Rosetta 2 (DE3) cells (EMD Millipore) was inoculated into 1,000 ml of LB medium containing ampicillin (50 μg/ml) and chloramphenicol (25 μg/ml) in a 4,000-ml Erlenmeyer flask and grown overnight with shaking at 37 °C. 50 ml of the starter culture was then added to 1,000 ml of LB medium containing ampicillin (50 μg/ml) in one to six 4,000-ml Erlenmeyer flasks and grown at 37 °C to O.D._600_ = 0.6-0.7, at which time protein expression was induced by adding 1 mM IPTG and incubating overnight with shaking at 19 °C. The cells were pelleted by centrifugation and stored at –80 °C overnight. After thawing on ice, the cells were lysed by sonication in 500 mM NaCl, 20 mM Tris-HCl pH 7.5, 20% glycerol, 1 mg/ml lysozyme, 0.2 mM phenylmethylsulfonyl fluoride (Roche), and the lysate was clarified by centrifugation at 30,000 x g for 60 min at 4 °C in a JA 25.50 rotor (Beckman). Nucleic acids in the clarified lysate were precipitated by slowly adding polyethyleneimine to the lysate with constant stirring in an ice bath to a final concentration of 0.4%, and then centrifuging at 30,000 x g for 25 min at 4 °C in a JA 25.50 rotor (Beckman). GsI-IIC RT and other cellular proteins were then precipitated from the supernatant with 60% saturating ammonium sulfate, pelleted at 30,000 x g for 25 min at 4 °C in a JA 25.50 rotor (Beckman), and resuspended in 25 mL A1 buffer (300 mM NaCl, 25 mM Tris-HCl pH 7.5, 10% glycerol). The protein mixture was then purified through two tandem 5-mL MBPTrap HP columns (GE Healthcare). After loading, the tandem column was washed with 5 column volumes (CVs) of A1 buffer, and the maltose-binding protein tagged GsI-IIC RT was eluted with 10 CVs of 500 mM NaCl, 25 mM Tris-HCl pH 7.5, 10% glycerol containing 10 mM maltose. The final fractions containing the RT were diluted to 200 mM NaCl, 20 mM Tris-HCl pH 7.5, 10% glycerol and loaded onto a 5 mL HiTrap Heparin HP column (GE Healthcare) and eluted with a 12 CV gradient from buffer A1 to A2 taking 0.5 mL fractions. Fractions containing GsI-IIC RT were identified by SDS-PAGE, pooled, and concentrated using an 30K centrifugation filter (Amicon). The concentrated protein was then dialyzed into 500 mM NaCl, 20 mM Tris-HCl pH 7.5, 50% glycerol, and 10 μl aliquots were flash frozen using liquid nitrogen and stored at −80°C. Thawed aliquots were only used once.

### DNA and RNA oligonucleotides

The DNA and RNA oligonucleotides used in this work are listed in Table S1. Most were purchased in RNase-free HPLC-purified form from Integrated DNA Technologies. Exceptions were the R2R primers and the cDNA mimic used in Fig. S3, which were 5’ end-labeled and purified by electrophoresis in a denaturing 8% polyacrylamide gel.

For biochemical assays, the R2R and primer extension (PE) primer [DNA primers](100 pmol) were labeled with [γ-^32^P]ATP (125 pmol; 6,000 Ci/mmol; 150 μCi/μl; Perkin Elmer) by incubating DNA with T4 polynucleotide kinase (10 units; New England Biolabs) for 30 min at 37 °C. A typical 10-μl reaction was then diluted to 40 μl with double-distilled H_2_O and extracted with an equal volume of phenol:chloroform:isoamylalcohol (25:24:1). Unincorporated nucleotides were removed from the aqueous phase by using a P-30 Microspin Column RNase Free (Bio-Rad), and the oligonucleotide concentration was measured by using the Qubit ssDNA Assay kit (Thermo Fisher Scientific). RNA acceptors used in Fig. S3 were labeled by the same method, and their concentrations were measured using a Qubit RNA HS Assay kit (Thermo Fisher Scientific). The concentration of unlabeled oligonucleotides was determined by spectrophotometry using a NanoDrop 1000 (Thermo Fisher Scientific).

### Template-switching and non-templated nucleotide addition (NTA) assays

Templateswitching reactions with GsI-IIC RT were carried out by using the previously described protocol with minor modifications (6). The RNA template/DNA primer starter duplex was based on that used for 3’-RNA-seq adapter addition in TGIRT-seq (Nottingham et al., 2016) and consists of a 34-nt RNA oligonucleotide containing an Illumina R2 sequence (R2 RNA) with a 3-blocking group (3SpC3; Integrated DNA Technologies; Table S1) annealed to a complementary DNA primer (R2R) that leaves either a blunt end, a single nucleotide 3’-DNA overhang end, or in some experiments, longer 3’-DNA overhang ends (Table S1). The oligonucleotides were annealed at a ratio of 1:1.2 to a yield a final duplex concentration of 250 nM by heating to 82 °C for 2 min and then slowly cooling to room temperature. Unless specified otherwise, reactions were done with 500 nM GsI-IIC RT, 50 nM R2 RNA/R2R DNA starter duplex, and 100 nM acceptor oligonucleotide in 25 μl of medium containing 200 mM NaCl, 5 mM MgCl2, 20 mM Tris-HCl pH 7.5, 5 mM fresh DTT, and an equimolar mix of 1 mM each dATP, dCTP, dGTP, and dTTP (Promega) to give 4 mM total dNTP concentration. Reactions were set up with all components except dNTPs and preincubated for 30 min at room temperature and then initiated by adding dNTPs. Reactions were incubated at 60 °C for times indicated in Figure Legends and stopped by adding 2.5-μl portions to 7.5 μl of 0.25 M EDTA. The products were further processed by adding 0.5 μl of 5 N NaOH and heating to 95 °C for 3 min to degrade RNA and remove tightly bound GsI-IIC RT, followed by a cooling to room temperature and neutralization with 0.5 μl of 5 N HCl. After adding formamide loading dye (5 μL; 95% formamide, 0.025% xylene cyanol, 0.025% bromophenol blue, 10 mM Tris-HCl pH 7.5, 6.25 mM EDTA), the products were denatured by heating to 99 °C for 10 min and placed on ice prior to electrophoresis in a denaturing 6% polyacrylamide gel containing 7 M urea, 89 mM Tris-borate and 2 mM EDTA, pH 8 at 65 W for 1.5 to 2 h. A 5’-end-labeled, single-stranded DNA ladder (10 to 200 nts: ss20 DNA Ladder, Simplex Sciences) was run in a parallel lane in most experiments. The gels were dried, exposed to an Imaging screen-K (Bio-Rad), and scanned using a Typhoon FLA 9500 (GE Healthcare). The Typhoon laser scanner was used for all other experiments where “phosphorimager” is indicated.

Non-templated nucleotide addition (NTA) reactions were done as described for template-switching reactions but in the absence of acceptor oligonucleotide.

### Analysis of template switching reactions using non-denaturing PAGE

The 50-nt acceptor RNA (100 pmol) was 5’-labeled with [γ-^32^P]ATP (125 pmol; 6,000 Ci/mmol; 150 μCi/μl; Perkin Elmer) by incubating with T4 polynucleotide kinase (10 units; New England Biolabs) for 30 min at 37 °C.

Template-switching reactions were done as described above using 5’-^32^P-labeled 50-nt acceptor RNA diluted with unlabeled 50-nt acceptor to reach the desired final concentration and 50 nM of starter duplex. Reactions were incubated at 60 °C for 15 min and stopped by adding 2.5-μL portions to 7.5 μL of 0.25 M EDTA and 1 μL of proteinase K (0.8 units, New England Biolabs) and incubating at room temperature for 30 min. After addition of 5 μL of 3x loading buffer (30% glycerol, 0.025% xylene cyanol, 0.025% bromophenol blue, 30 mM Tris, 150 mM Hepes, pH 7.5), 5 μL of the sample was loaded onto a 8% polyacrylamide gel containing 34 mM Tris, 66 mM Hepes, pH 7.5, 0.1 mM EDTA and 3 mM MgC_2_ The gel was then run at 4°C for 4 h at a constant power of 3 W and then dried and imaged as described above for denaturing polyacrylamide gels. The cDNA mimic oligonucleotide used in the native PAGE experiment (Fig. S3) consists of the R2R-G sequence fused to the reverse complement of the 50-nt acceptor lacking the terminal C residue. The cDNA mimic was annealed to R2 RNA and 50-nt acceptor C oligonucleotides to provide a marker for template-switching product bands in the gel.

### Primer extension assays

Primer extension reactions with GsI-IIC RT were carried out similarly to template-switching reactions by using a 50-nt, 5’-end-labeled DNA primer (PE primer) annealed near the 3’ end of a 1.1-kb *in vitro* transcribed RNA. The transcript was generated by T3 runoff transcription (T3 MEGAscript kit, Thermo Fisher Scientific) of pBluescript KS (+) (Agilent) linearized using XmnI (New England Biolabs) and cleaned up using a MEGAclear kit (Thermo Fisher Scientific). The labeled DNA primer was annealed to the RNA template at a ratio of 1:1.2 to a yield a final duplex concentration of 250 nM by heating to 82 °C for 2 min followed by slowly cooling to room temperature. GsI-IIC RT (500 nM) was pre-incubated with 50 nM of the annealed template-primer in 25 μl of reaction medium containing 200 mM NaCl, 5 mM MgCl2, 20 mM Tris-HCl pH 7.5, 5 mM DTT for 30 min at room temperature, and reverse transcription was initiated by adding 1 μl of the 25 mM dNTP mix to give a final dNTP concentration of 1 mM for each dNTP. After incubating at 60 °C for 15 min, the reaction was terminated, processed, and analyzed by electrophoresis in a denaturing 8% polyacrylamide gel, as described above for template-switching reactions.

### RNA Sequencing

RNA sequencing libraries were prepared as described (8, 9) by using TGIRT-III (InGex), a commercial version of GsI-IIC RT, with different R2 RNA/R2R starter duplexes and acceptor nucleic acids as indicated in the text. The initial template template-switching reactions for addition of the R2R adapter to the 5’ end of the cDNA were done as described above with 500 nM TGIRT-III enzyme, 50 nM unlabeled starter duplex, and 100 nM acceptor RNA for 15 min at 60 °C. After terminating the reactions with NaOH and neutralization with HCl as described above for template-switching reactions, the volume was raised to 100 μl with H_2_O, and cDNA products containing the R2R adapter attached to their 5’ end were cleaned-up by using a MinElute column (Qiagen) to remove unused primer. A 5’ adenylated R1R adapter was then ligated to the 3’ end of the cDNA using Thermostable 5’ AppDNA/RNA ligase (New England Biolabs) for 1 h at 65 °C. After another MinElute clean up, the entirety of the eluent was used for a 12 cycle PCR reaction using Phusion polymerase (ThermoFisher), and the resulting libraries were cleaned up by using 1.4x Ampure XP beads to remove residual primers, primer dimers, salts, and enzymes. The quality of the libraries was assessed by using a 2100 Bioanalzyer Instrument with a High Sensitivity DNA chip (Agilent). The libraries were sequenced on an Illumina NextSeq instrument to obtain 1-2 million 75-nt paired-end reads. Read 1 was used for analysis.

To analyze template-switching junctions, the 75-nt-reads 1 from each dataset were trimmed from the 5’ end using Cutadapt v2.5 (64) to remove all but the last 3 nucleotides (GAC) of the adapter proximal acceptor by using the following parameters: cutadapt −O 10 --nextseq-trim=20 --trim-n −q 20 --discard-untrimmed −g CGCCGGACCGTGCACCATCTGGAGTTATAGAGATGAGTCTCACATA -j 8 -e 0.1 -o {Step1 trimmed reads} {Read 1 file}. The reads were then trimmed from the 3’ end to leave the junction sequences flanked by GAC at the 5’ end and CGC (acceptor) or AGA (donor) at the 3’ end by using the following parameters: cutadapt -O 10 --nextseq-trim=20 --trim-n -q 20 --discard-untrimmed -a CGGACCGTGCACCAT -a TCGGAAGAGCACACG -j 8 -e 0.1 -o {Step2 trimmed reads} {Step1 trimmed reads}. Finally, the trimmed reads containing either the acceptor-donor (5’-GAC-(N)n-AGA-3’) or acceptor-acceptor (5’-GAC-(N)n-CGC-3’) junction were binned and counted by using lab scripts to determine the frequencies of different junctions.

### Analysis of kinetics experiments

Phosphorimager scans of reaction time courses were quantified with ImageQuant TL 8.1 (GE Healthcare) by generating rectangular boxes around the unextended primer and each reaction product. Background was subtracted by using analogous rectangles on a portion of the screen that did not to correspond to a gel lane. Fractions of product were plotted versus time and fit by a single-exponential function. For analyses of concentration dependence, rate constants were plotted against the concentration of the species being varied (enzyme or dNTP) and fit by a hyperbolic equation to obtain values of the maximal rate constant and half-maximal concentration of the varied species. Data fitting was done using Prism 8 (GraphPad). Unless otherwise indicated, the reported uncertainty values reflect the standard error obtained from these fits. For reactions that were performed in triplicate (*e.g.*, Fig. 1B), fitting each data set individually gave rate constants with standard error values after averaging of <25% and amplitudes with standard error values after averaging of <5% (not shown), reflecting the level of day-to-day variability in these reactions.

Reactions monitoring NTA of individual nucleotides were analyzed by global modeling using Kintek Explorer (45). Data were fit by a model that included up to three equilibrium binding steps, three irreversible extensions, and paused and stopped states at each cycle. To obtain an initial fit, we set the reaction rates and equilibrium binding constants to those derived from conventional fitting of the data to analytic functions. The fit was further refined by adjusting the reaction rates to approximate the data and then using the data fit editor to fit globally to data from reactions at four different dNTP concentrations. This process was repeated iteratively to obtain the final fits. Plots of normalized chi^2^ values versus the rate constant parameter (1D Fitspace) are shown in Fig. S6.

## Acknowledgements

*This work was supported by National Institutes of Health Grants R01 GM37949 to A.M.L. and R35 GM131777 to R.R. The content is solely the responsibility of the authors and does not necessarily represent the official views of the National Institutes of Health. We thank Dr. Jennifer Stamos for comments on the manuscript. RNA sequencing data have been deposited in the Gene Expression Omnibus (GEO) under accession number GSE138200.*

## Conflict-of-interest statement

*Thermostable group II intron reverse transcriptase enzymes and methods for their use are the subject of patents and patent applications that have been licensed by the University of Texas and East Tennessee State University to InGex, LLC. A.M.L, some former and present members of AML’s laboratory, and the University of Texas are minority equity holders in InGex, LLC and receive royalty payments from the sale of TGIRT enzymes and kits employing TGIRT template-switching activity for RNA-seq adapter addition and from the sublicensing of intellectual property to other companies.*

## Footnotes

*The abbreviations used are:* 2’-*O*-Me, 2’-*O*-Methyl; cDNA, complementary DNA; CV, column volume; LINE, long interspersed nuclear element; LTR, long terminal repeat; NTA, non-templated nucleotide addition; NTE, N-terminal extension; MRP, mitochondrial retroplasmid; mt, mitochondrial; RdRP, RNA-dependent RNA polymerase; RT, reverse transcriptase; SMART-Seq, switching mechanism at the 5’ end of the RNA transcript sequencing; spp., species; TGIRT, thermostable group II intron RT; TGIRT-seq, RNA-seq using thermostable group II intron RT; TPRT, target DNA-primed reverse transcription.

---

**FIGURE S1.**
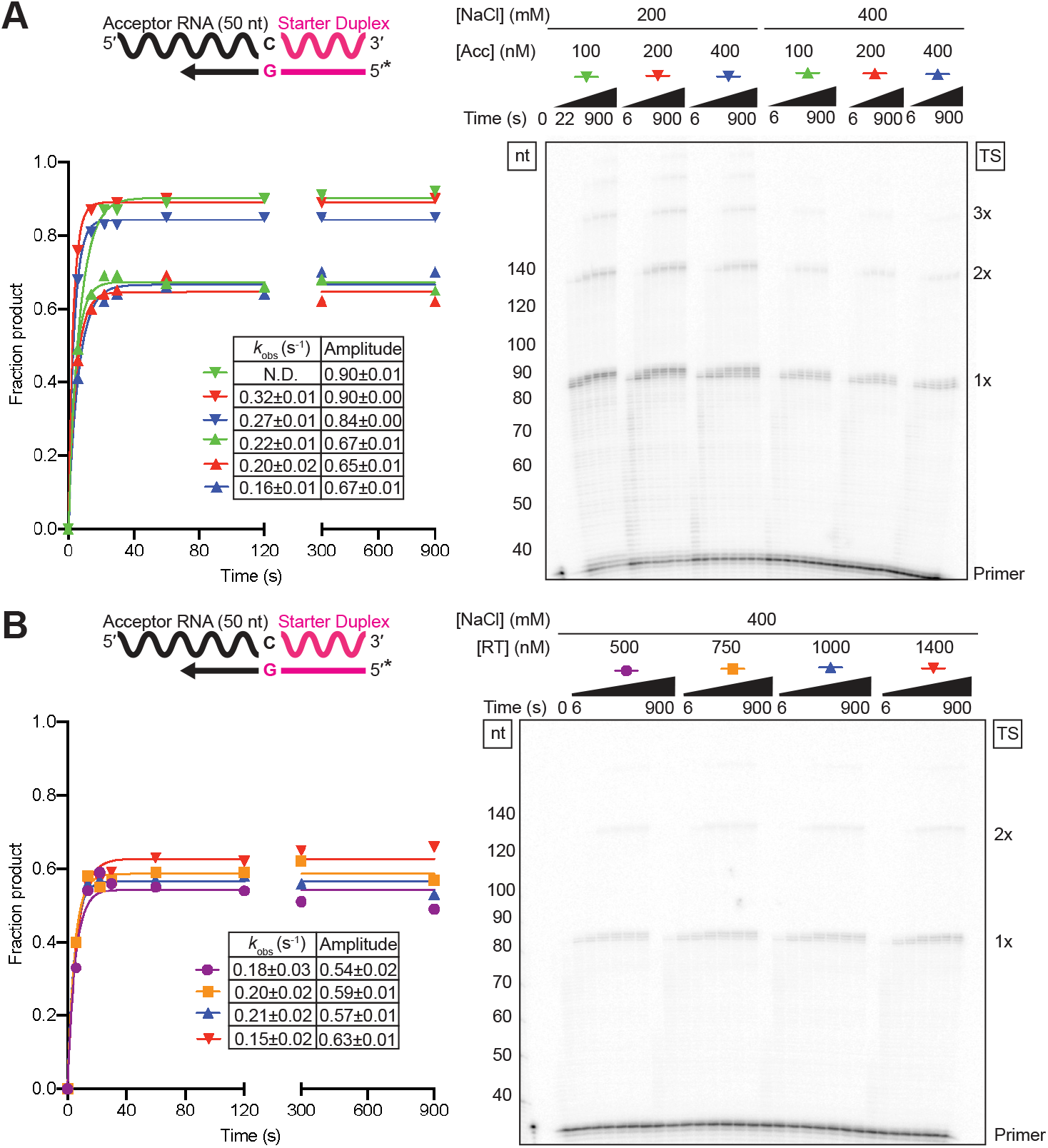
Higher concentrations of GsI-IIC RT or acceptor RNA do not increase amplitudes in template-switching reactions at high salt concentration. *A*, Template-switching reactions with 500 nM GsI-IIC RT and varying concentrations of 50-nt acceptor RNA in reaction media containing 200 or 400 mM NaCl. *B*, Template-switching reactions with 100 nM of 50-nt acceptor RNA and varying concentrations of GsI-IIC RT in reaction medium containing 400 mM NaCl. Template-switching reactions were done and analyzed and the gels were labeled as in Fig. 1.

**FIGURE S2.**
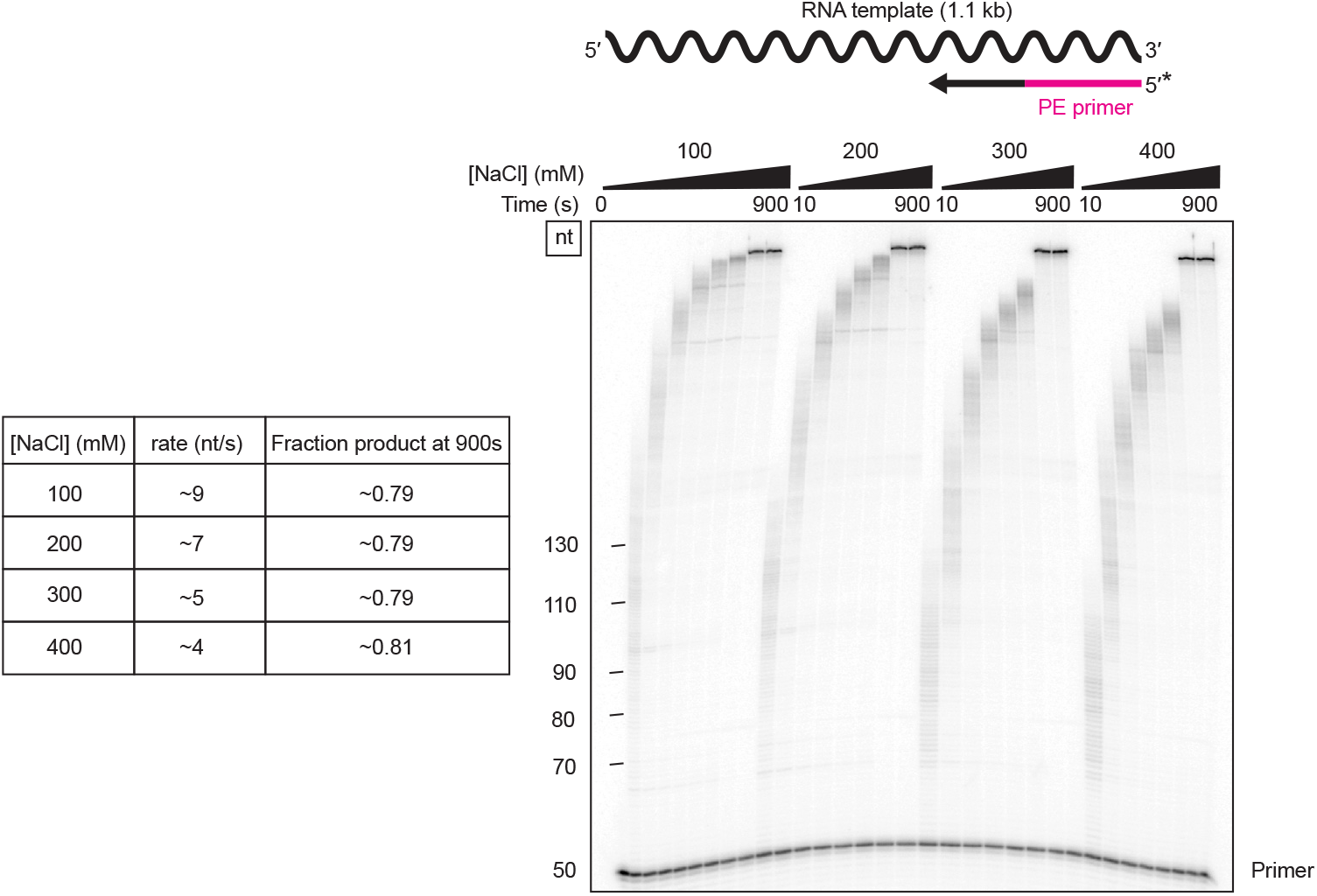
High salt concentrations have relatively little effect on primer extension by GsI-IIC RT. Primer extension reactions were done with 50 nM template-primer substrate (a 1.1-kb *in vitro* transcribed RNA with a 50-nt ^32^P-labeled DNA primer (PE primer; Table S1) annealed at its 3’ end) and 500 nM GsI-IIC RT in reaction media containing varying NaCl concentrations. The reaction was stopped after times ranging from 10 to 900 s, and the products were analyzed by denaturing PAGE and quantified from a phosphorimager scan of the dried gel. cDNA synthesis rates were approximated by the mean progression of nucleotide bands. The fraction product was measured by quantifying products larger than the primer and remained largely unchanged throughout the reaction. Extension rates and the end point fraction product are shown to the left.

**FIGURE S3.**
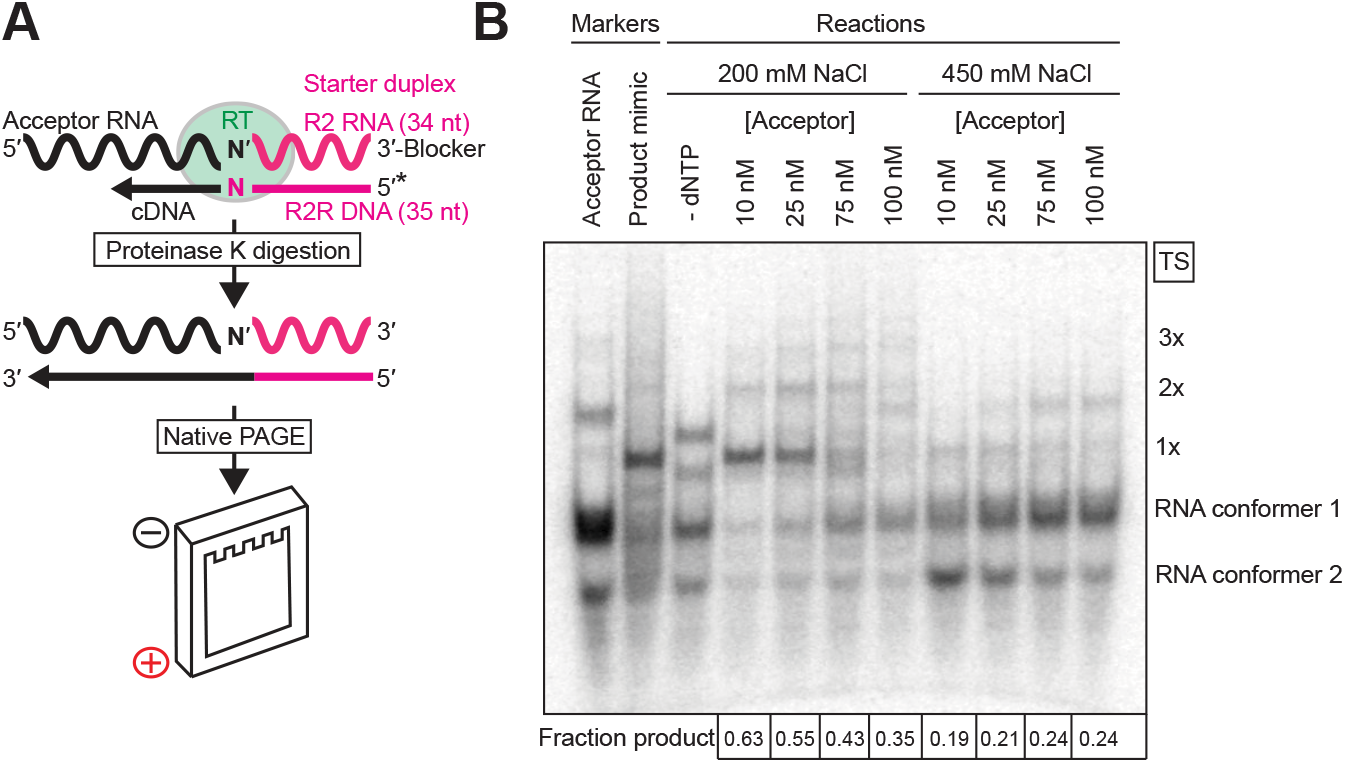
Template-switching reactions with varying concentrations of a 5’-labeled acceptor RNA analyzed by nondenaturing PAGE. Template-switching reactions with varying concentrations of 50-nt acceptor RNA with a 3’-C residue and 50 nM of starter duplex with a 1-nt 3’-G overhang were done for 15 min at 60 °C in reaction medium containing 200 or 450 mM NaCl. After treatment with proteinase K to remove GsI-IIC RT, the products were analyzed by electrophoresis in a nondenaturing 8% polyacrylamide gel. The acceptor RNA (denatured by heating to 82 °C in 1x TE and placed on ice), and a product mimic (labeled acceptor RNA and R2 RNA annealed to a complementary 84-nt oligonucleotide corresponding to the sequence of a full-length cDNA) were loaded in parallel lanes. The gel was run at 4 °C at a constant power of 3 W to maintain the duplex structure and then dried and scanned with a phosphorimager. The labels to the right of the gel indicate the products resulting from the initial template switch (1x) and subsequent end-to-end template switches from the 5’ end of one acceptor to the 3’ end of another (2x, 3x, *etc*.). Putative product bands were summed and divided by the product and substrate bands to obtain a measure of the fraction product formed, which is indicated under each lane.

**FIGURE S4.**
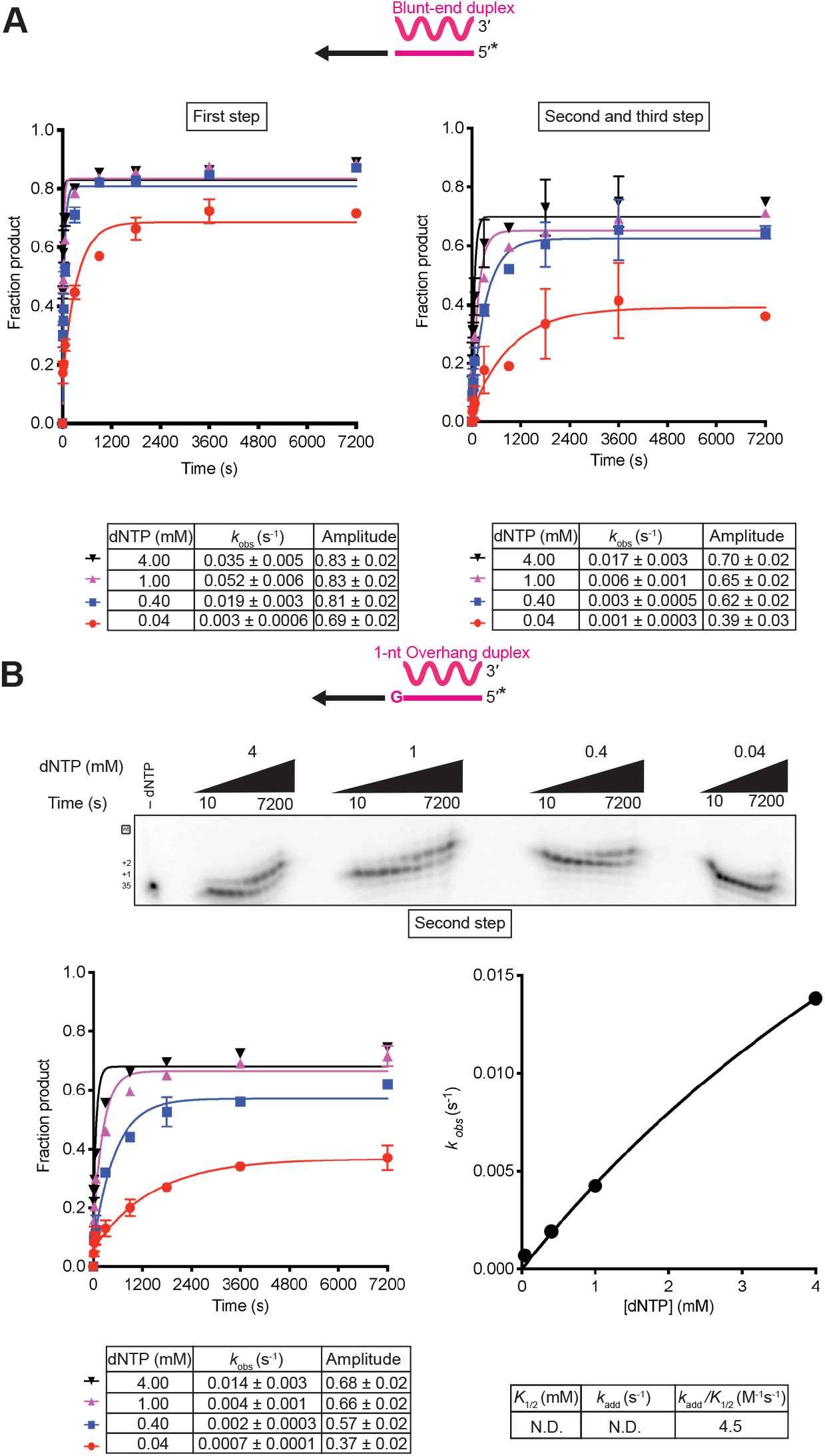
Non-templated nucleotide addition activity to blunt-end and 1-nt 3’ overhang starter duplexes using mixed dNTPs. *A*, Reactions included 500 nM GsI-II RT and 50 nM of a blunt-end starter duplex with 5’-^32^P-labeled (*) DNA primer in a solution containing 200 mM NaCl and varying dNTP concentrations (0.04, 0.4, 1, and 4 mM, where 4 mM is an equimolar mix of 1 mM dATP, dCTP, dGTP, and dTTP). Aliquots were quenched at times from 10 s to 7,200 s, and the products were analyzed by electrophoresis in a denaturing polyacrylamide gel, which was dried and scanned with a phosphorimager (gels and calculated kinetic parameters shown in Fig. 4). *B*, Reactions included 500 nM GsI-II RT and 50 nM of a 1-nt G overhang starter duplex and 200 mM NaCl. Reaction aliquots were stopped at times from 10 to 7,200 s, and the products were analyzed as in A. The observed rates for the first NTA from this template were used to calculate the kinetic parameters shown below the plot.

**FIGURE S5.**
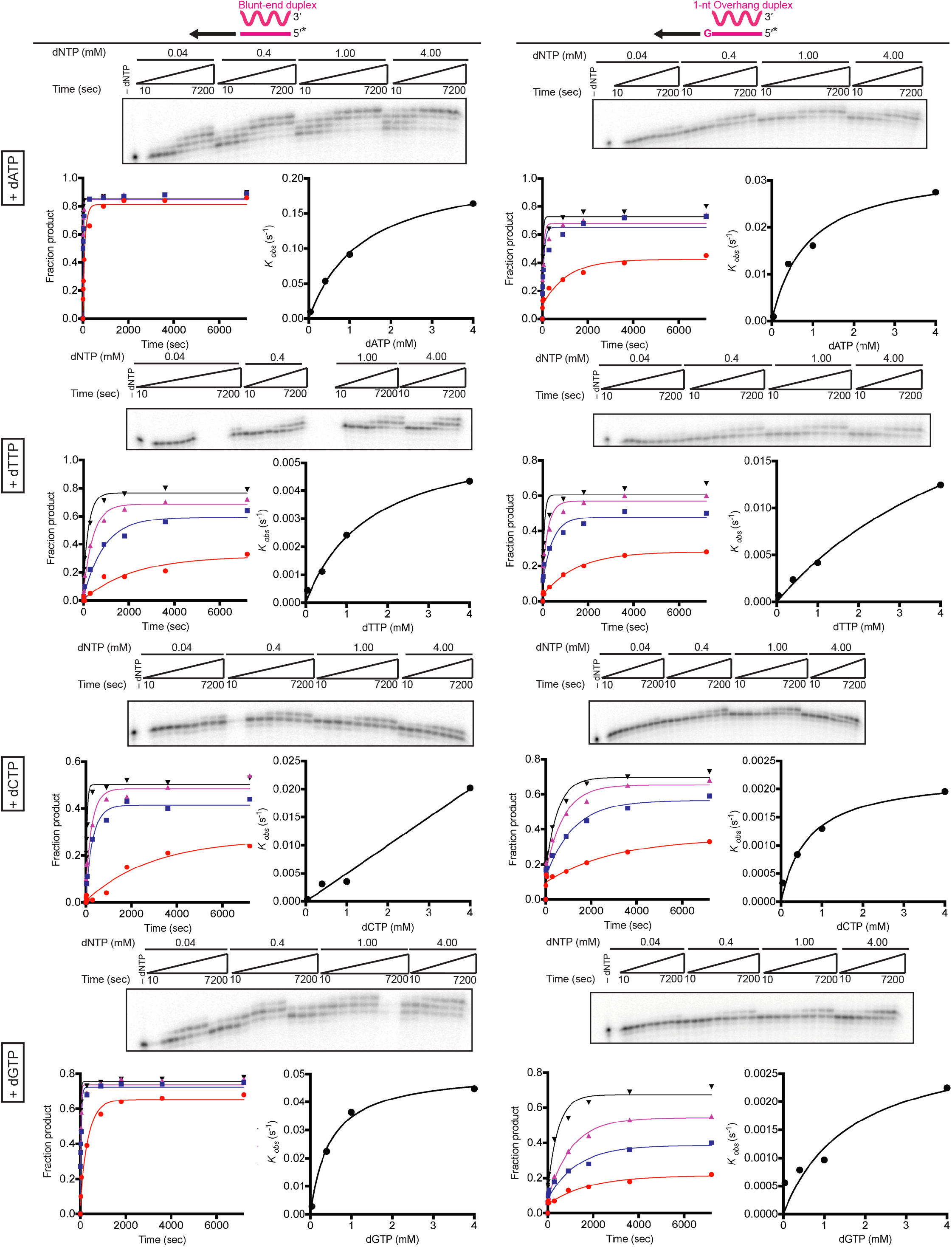
Non-templated nucleotide addition reactions to blunt-end and 1-nt 3’ overhang starter duplexes using individual dNTPs. *A*, Assays of non-templated nucleotide addition to the ^32^P-labeled blunt-end RNA/DNA duplex (left) and 1-nt 3’-overhang RNA/DNA duplex (right) using 0.04, 0.4, 1, and 4 mM dATP, dTTP, dCTP and dGTP individually. Reactions included 500 nM GsI-IIC RT and 50 nM of starter duplex in reaction medium containing 200 mM NaCl and were stopped at times ranging from 10 to 7,200 s. The reaction products were analyzed as in Fig. S4. The values obtained from these experiments were used for global fitting analysis in Fig. S6 and for the summary of second-order rate constants in Fig. 5.

**FIGURE S6.**
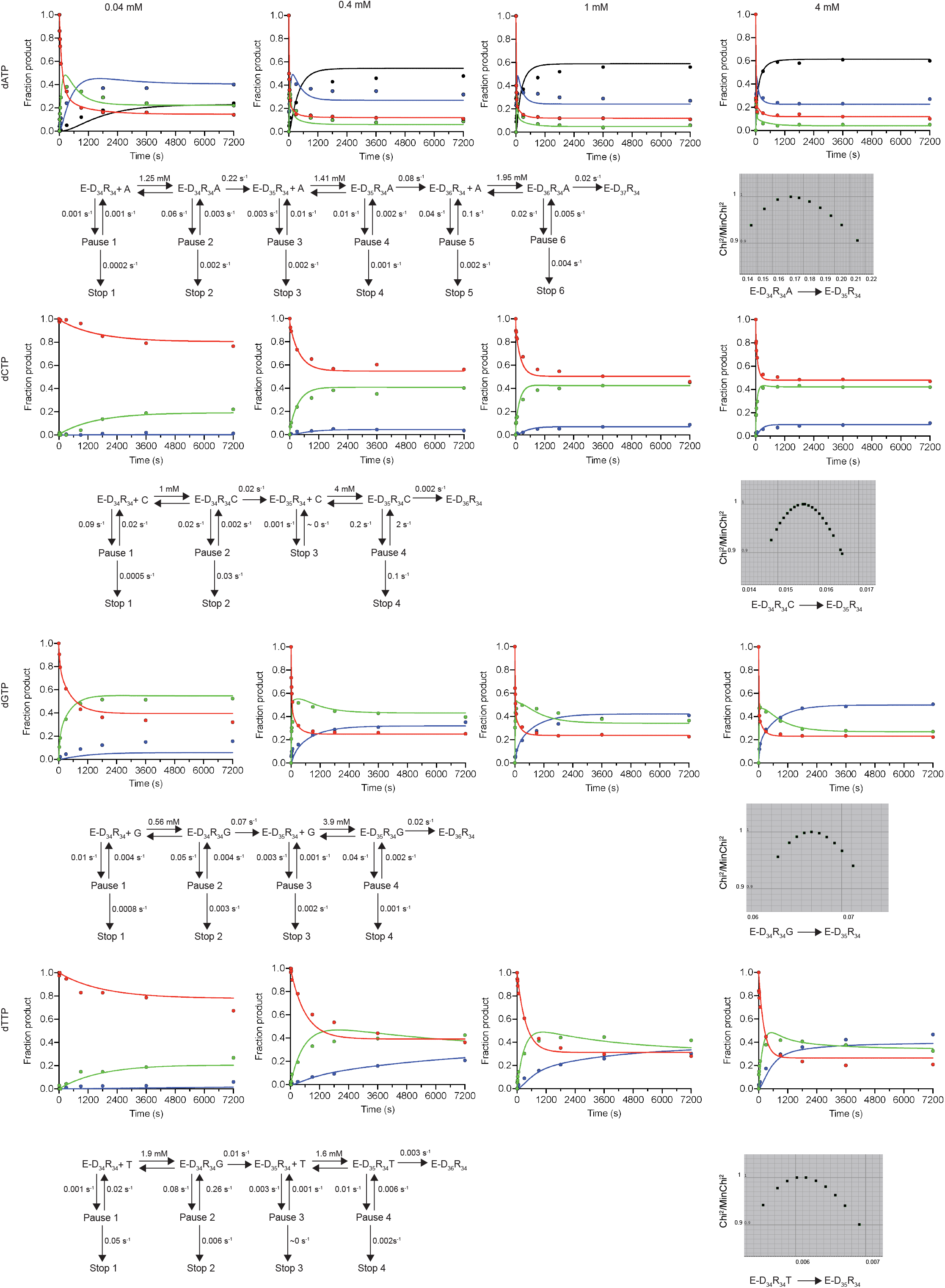
Global fitting of consecutive non-templated nucleotide addition using individual dNTPs. Data and resulting models are shown for NTA of individual dNTPs. For dATP, three NTA steps were observed, and for the other dNTPs only two steps were observed. The plots show global data fitting of consecutive dNTP addition at the indicated concentrations of each dNTP. Each color represents a unique species in the reaction pathway: red, blunt-end starter duplex; green, product after first NTA; blue, product after second NTA; black, product after third NTA (if detected). For details on the global fitting process, see Experimental Procedures. A 1D fitspace plot is shown for each dNTP for the first reaction of nucleotide addition. These plots show the normalized chi^2^ values for a range of values for the rate constant, with a chi^2^ threshold of 0.9.

**FIGURE S7.**
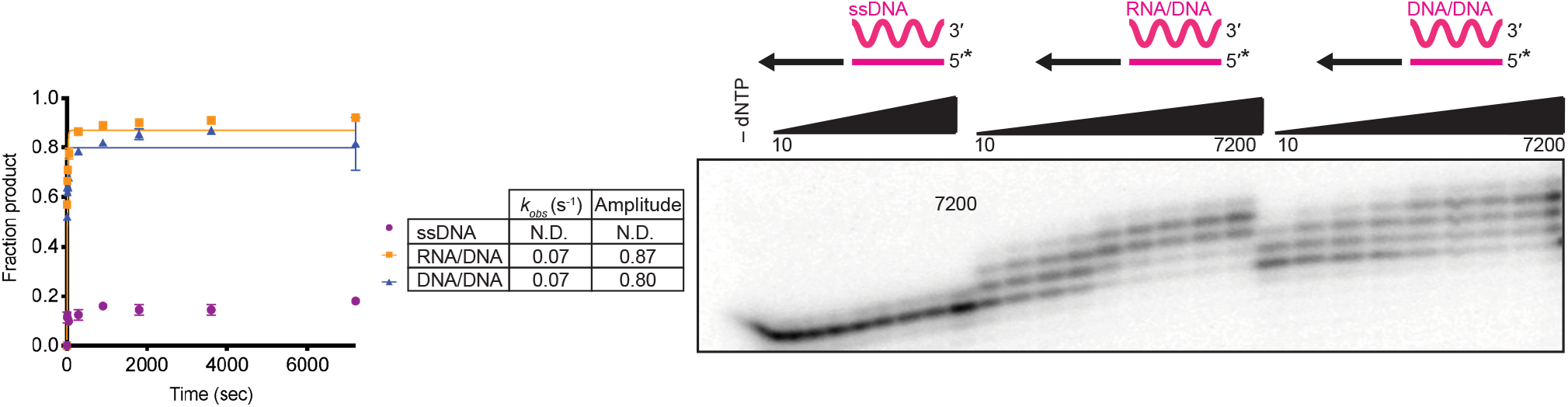
Non-templated addition reactions of various starter molecule conformations. Assays of NTA to 5’-^32^P-labeled (*) single-stranded (ss) DNA (34 nt R2R DNA; left), a 34-bp RNA/DNA duplex (R2 RNA/R2R DNA; center), and a 34-bp DNA/DNA duplex (R2 DNA/R2R DNA; right). Reactions were done with 500 nM GsI-IIC RT and 50 nM substrate in reaction medium containing 200 mM NaCl and 4 mM dNTPs (an equimolar mix of 1 mM dATP, dCTP, dGTP, and dTTP) with time points ranging from 10 to 7,200 s. The reaction products were analyzed as in Fig. S4.

**FIGURE S8.**
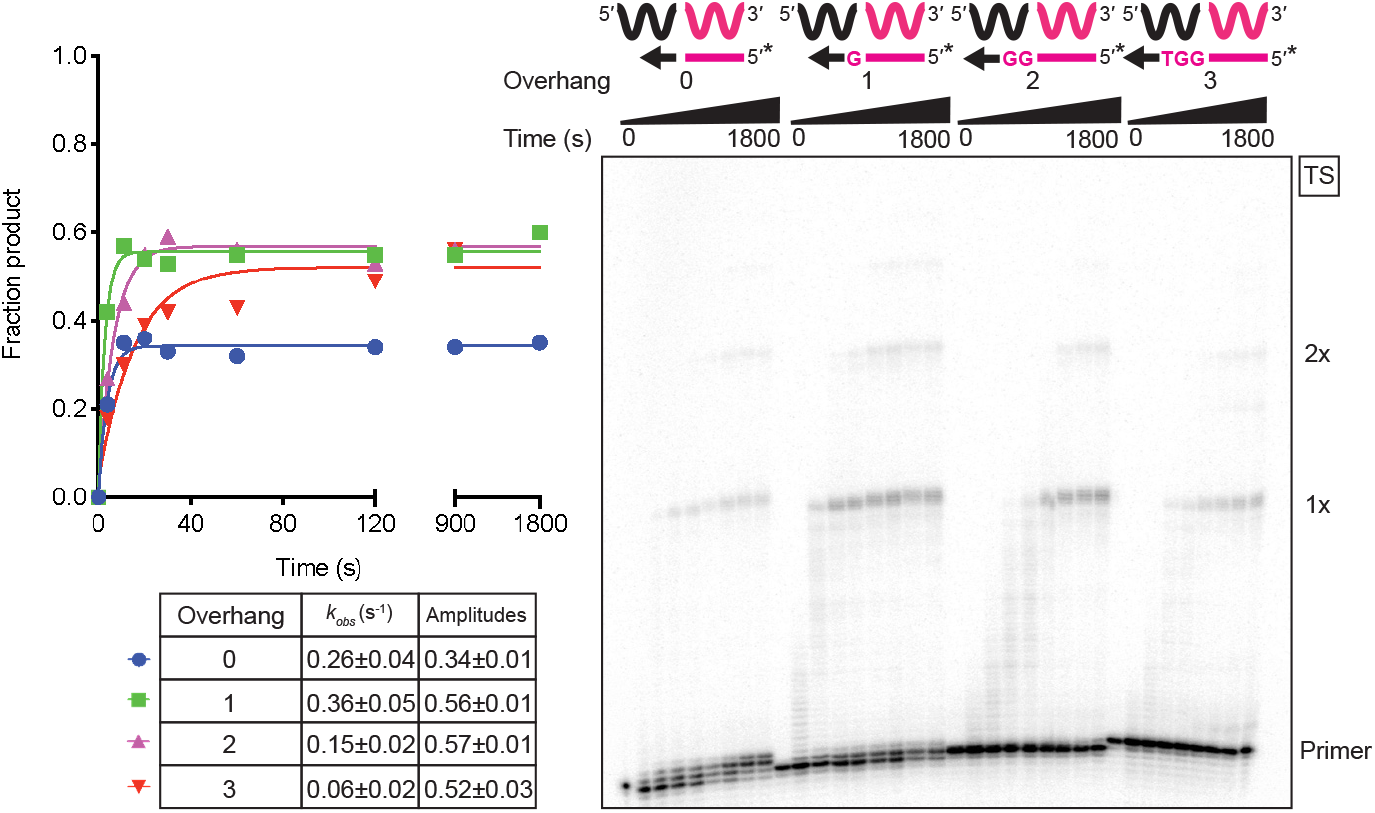
Template switching to DNA acceptors is favored by a single nucleotide 3’ overhang. Template-switching reactions from starter duplexes with a blunt end (0 overhang) or 1, 2, or 3-nt 3’ overhangs were done with 500 mM GsI-IIC RT, 100 nM 50-nt acceptor DNA, and 50 nM starter duplex with a ^32^P-labeled (*) DNA primer in reaction medium containing 200 mM NaCl at 60 ^o^C. Reactions were stopped after times ranging from 5 to 1,800 s, and the products were analyzed by denaturing PAGE and quantified from phosphorimager scans of the dried gel. The plots to the left of the gel show the data fit by a single exponential function to calculate the *k*_obs_ and amplitude for each time course, and the values are summarized in the table below the plots together with the standard error of the fit. The gel is labeled as in Fig. 1.

**FIGURE S9.**
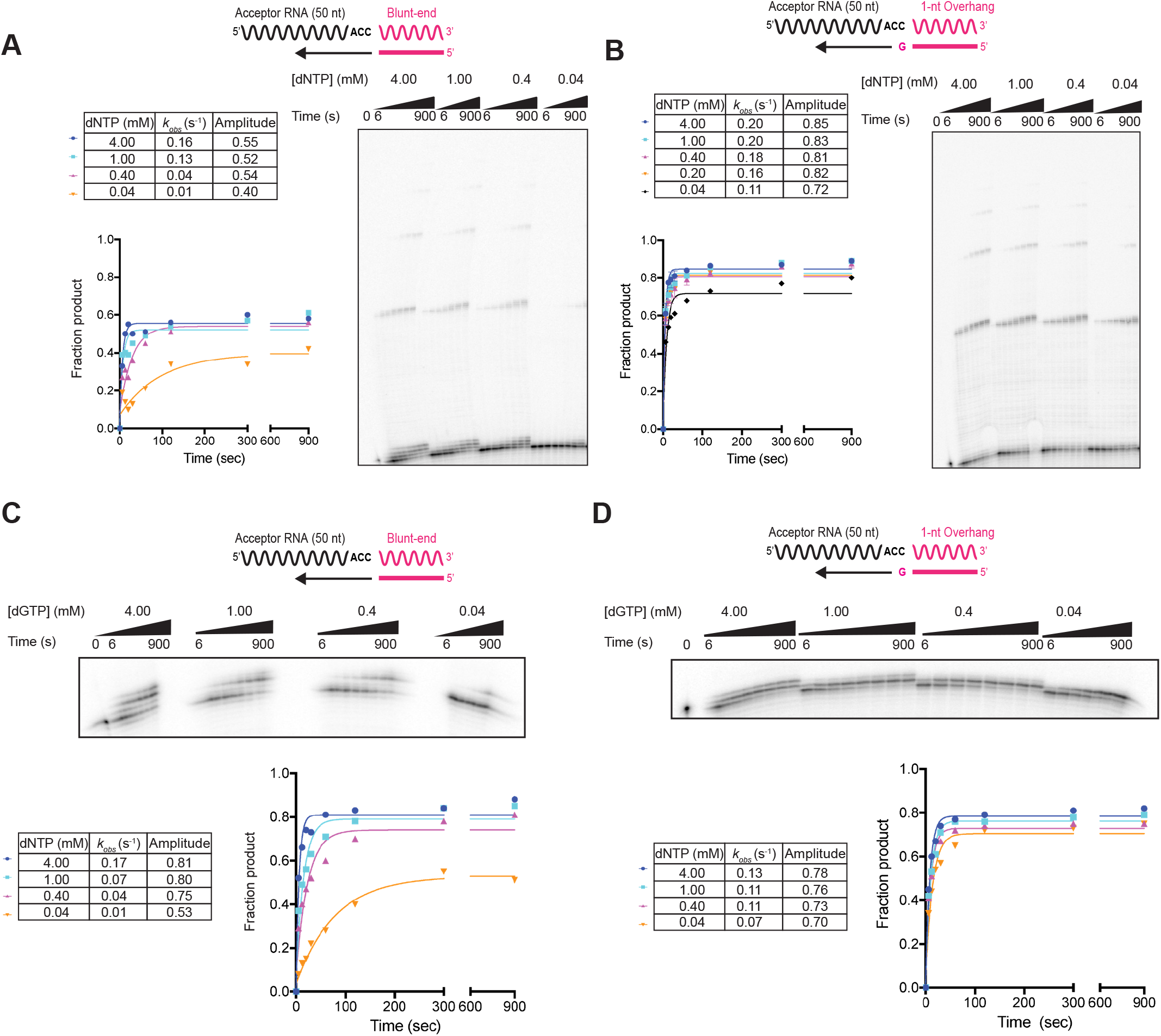
Template-switching reactions from blunt-end and 1-nt 3’-overhang starter duplexes with varying concentrations of dNTPs. *A* and *B*, Template-switching reactions with 50 nM blunt-end (A) or 1-nt overhang (B) starter duplexes with a ^32^P-labeled (*) DNA primer and 100 nM 50-nt acceptor RNA were done with 500 nM GsI-IIC RNA RT and 0.04, 0.4, 1 or 4 mM dNTPs (where 4 mM dNTPs = 1 mM each of dATP, dCTP, dGTP, and dTTP). The reactions were stopped after times ranging from 6 to 900 s, and the products were analyzed by denaturing PAGE and quantified from a phosphorimager scan of the dried gel. The data were fit by a single exponential function to obtain the observed rates and amplitudes, which are summarized in the associated tables. *C* and *D*, The same template-switching reactions as in A and B but adding 0.04, 0.4, 1, or 4 mM dGTP, the nucleotide complementary to the last 2 nucleotides (CC) of the acceptor RNA, so that the template-switching products for the (C) blunt-end duplex is 2 nt longer than the DNA primer, and that for the (D) 1-nt overhang duplex is only 1 nt longer than the DNA primer.

**FIGURE S10.**
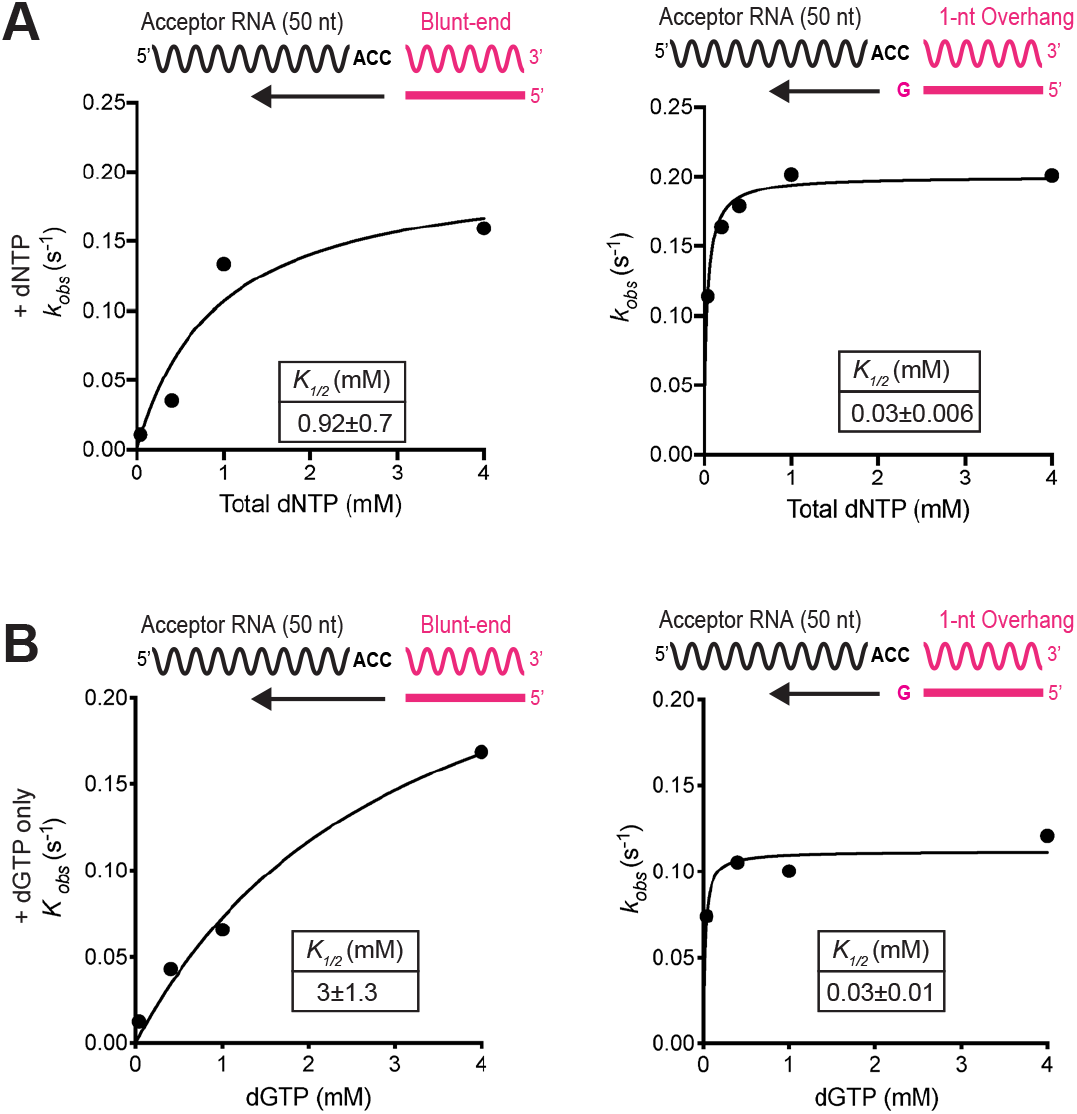
Template switching from a blunt-end duplex has a higher *K*_1/2_ value for dNTPs than does template switching from a 1-nt 3’ overhang duplex. *A and* B, the observed rates for template switching at varying (A) dNTP or (B) dGTP concentrations from Fig. S9 were plotted against dNTP concentration and fit by a hyperbolic function to obtain *K*_1/2_.

**TABLE S1.**
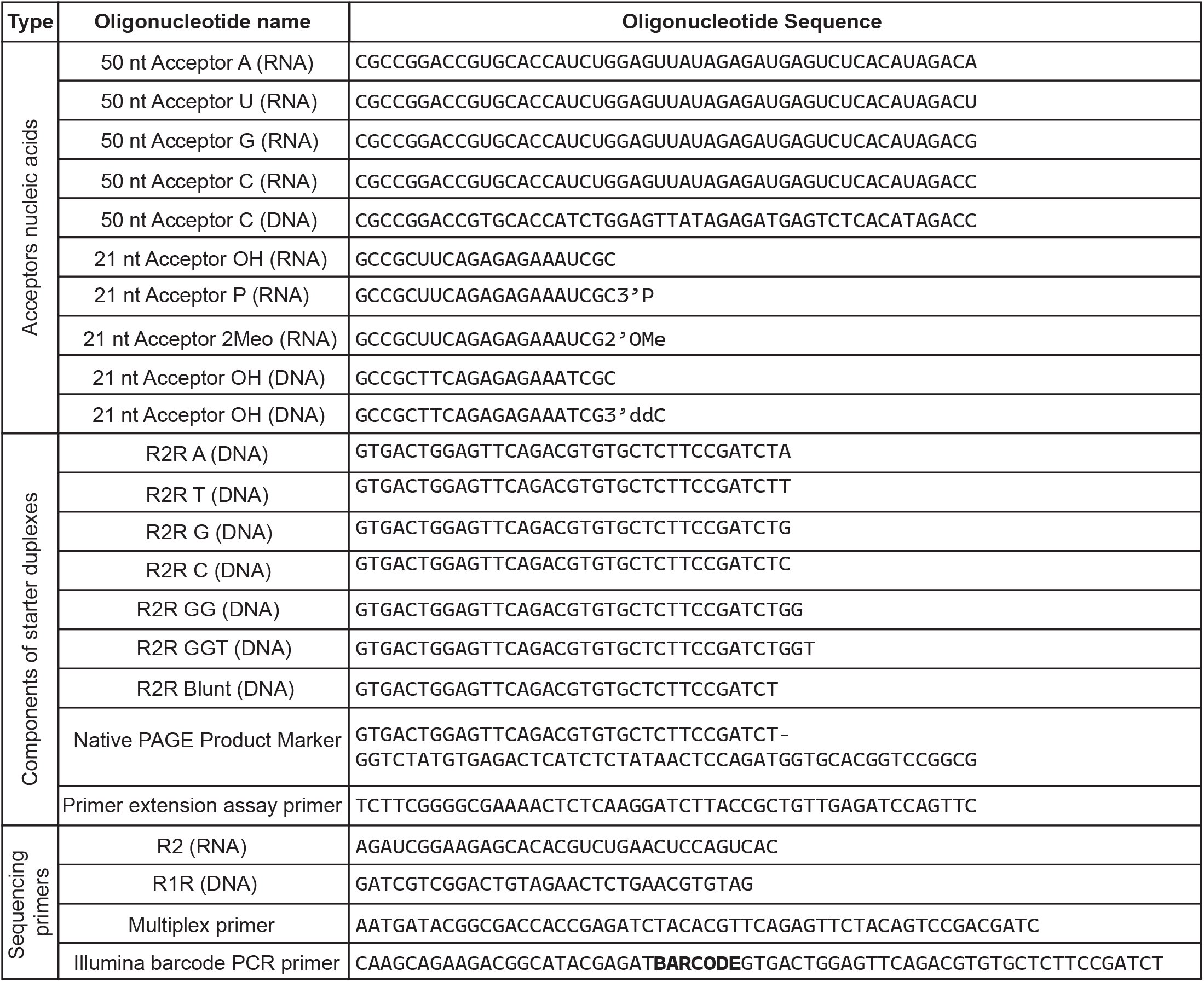
Sequences of oligonucleotides used in this study. The oligonucleotides are divided into three sections indicated in the left-hand column: acceptor nucleic acids, components of starter duplexes, and sequencing primers.

**TABLE S2.** Kinetic parameters for non-templated nucleotide addition of individual nucleotides. The kinetic parameters of non-templated nucleotide addition with individual nucleotides, with the data shown in Figure S5 and second order rate constants in Fig. 5A.

| Duplex type | dNTP | dNTP (mM) | $k_{obs}$ (s<sup>-1</sup>) | Product amplitude | $K_{1/2}$ (mM) | $k_{add}$ (s<sup>-1</sup>) | $k_{add}/K_{1/2}$ (M<sup>-1</sup>s<sup>-1</sup>) |
| --- | --- | --- | --- | --- | --- | --- | --- |
| Blunt | dATP | 4.00 | 0.16 ± 0.03 | 0.85 ± 0.02 | 1.3 ± 0.14 | 0.22 ± 0.01 | 170 |
|  |  | 1.00 | 0.09 ± 0.02 | 0.85 ± 0.02 |  |  |  |
|  |  | 0.40 | 0.05 ± 0.01 | 0.85 ± 0.03 |  |  |  |
|  |  | 0.04 | 0.01 ± 0.002 | 0.81 ± 0.03 |  |  |  |
|  | dTTP | 4.00 | 0.004 ± 0.001 | 0.76 ± 0.03 | 1.6 ± 0.4 | 0.006 ± 0.0006 | 4 |
|  |  | 1.00 | 0.002 ± 0.0004 | 0.69 ± 0.02 |  |  |  |
|  |  | 0.40 | 0.001 ± 0.0002 | 0.59 ± 0.03 |  |  |  |
|  |  | 0.04 | 0.0004 ± 0.0002 | 0.32 ± 0.04 |  |  |  |
|  | dCTP | 4.00 | 0.02 ± 0.005 | 0.50 ± 0.02 | N.D. | N.D. | 9 |
|  |  | 1.00 | 0.004 ± 0.001 | 0.49 ± 0.02 |  |  |  |
|  |  | 0.40 | 0.003 ± 0.0007 | 0.41 ± 0.02 |  |  |  |
|  |  | 0.04 | 0.0004 ± 0.0001 | 0.27 ± 0.04 |  |  |  |
|  | dGTP | 4.00 | 0.04 ± 0.006 | 0.75 ± 0.02 | 0.5 ± 0.08 | 0.05 ± 0.003 | 100 |
|  |  | 1.00 | 0.04 ± 0.007 | 0.74 ± 0.03 |  |  |  |
|  |  | 0.40 | 0.02 ± 0.004 | 0.72 ± 0.03 |  |  |  |
|  |  | 0.04 | 0.003 ± 0.0006 | 0.65 ± 0.02 |  |  |  |
| Overhang | dATP | 4.00 | 0.03 ± 0.007 | 0.73 ± 0.03 | 0.9 ± 0.25 | 0.03 ± 0.003 | 33 |
|  |  | 1.00 | 0.02 ± 0.005 | 0.68 ± 0.04 |  |  |  |
|  |  | 0.40 | 0.01 ± 0.005 | 0.65 ± 0.04 |  |  |  |
|  |  | 0.04 | 0.001 ± 0.0004 | 0.42 ± 0.04 |  |  |  |
|  | dTTP | 4.00 | 0.012 ± 0.005 | 0.60 ± 0.03 | 6.0 ± 2.4 | 0.03 ± 0.008 | 5 |
|  |  | 1.00 | 0.004 ± 0.002 | 0.57 ± 0.03 |  |  |  |
|  |  | 0.40 | 0.002 ± 0.0009 | 0.48 ± 0.03 |  |  |  |
|  |  | 0.04 | 0.0007 ± 0.0001 | 0.28 ± 0.01 |  |  |  |
|  | dCTP | 4.00 | 0.002 ± 0.0007 | 0.70 ± 0.04 | 0.7 ± 0.23 | 0.002 ± 0.0003 | 3 |
|  |  | 1.00 | 0.001 ± 0.0005 | 0.65 ± 0.05 |  |  |  |
|  |  | 0.40 | 0.0008 ± 0.0003 | 0.57 ± 0.05 |  |  |  |
|  |  | 0.04 | 0.0003 ± 0.0002 | 0.35 ± 0.08 |  |  |  |
|  | dGTP | 4.00 | 0.002 ± 0.0006 | 0.67 ± 0.04 | 1.6 ± 1.5 | 0.003 ± 0.001 | 2 |
|  |  | 1.00 | 0.001 ± 0.0002 | 0.54 ± 0.03 |  |  |  |
|  |  | 0.40 | 0.0007 ± 0.0003 | 0.38 ± 0.04 |  |  |  |
|  |  | 0.04 | 0.0006 ± 0.0002 | 0.21 ± 0.03 |  |  |  |

TABLE S3. **RNA-seq analysis of template-switching junctions**.

